# Forest belowground productivity and carbon allocation predominantly driven by soil properties rather than climate

**DOI:** 10.64898/2026.09.24.754289

**Authors:** Anne Y. Polyakov, Angela M. Klock, Korena K. Mafune, Andrew M. Berdahl, Kristiina A. Vogt, Daniel J. Vogt

## Abstract

Forests are threatened by a multitude of stressors, including anthropogenic disturbances and climate change. Assessing how forests will respond to these stressors requires a comprehensive understanding of net primary productivity (Npp), environmental constraints on growth, and adaptive capacity. A parameter of significant uncertainty is belowground Npp (bNpp), which can account for up to 80% of total Npp but is poorly estimated and rarely measured directly. We used a cross-biome dataset of direct, field-based measurements of aboveground and belowground primary productivity and 21 climatic and soil variables to identify potential constraints on bNpp and belowground carbon allocation in boreal and cold temperate forests. Soil variables, rather than climate variables, were the main drivers of bNpp and belowground allocation across biomes. The importance of soil variables suggests that soil nutrient dynamics, especially soil nutrient pool and flux variables, must be explicitly modeled to more accurately predict feedbacks between climate, productivity, and within-tree carbon allocation. Within biomes, environmental drivers of belowground allocation varied between low versus high allocation forests, indicating that environmental drivers are site-specific and the development of within-biome, site-scale classifications for forest ecosystems could be useful. Changes in soil variables, such as increasing soil nitrogen pools, caused abrupt and large decreases in bNpp for boreal, but not cold temperate forests. Threshold-like shifts indicate that boreal forests might have lower adaptive capacity and higher sensitivity to disturbances than cold temperate forests. With 70% of boreal forests characterized by low bNpp, disturbances such as anthropogenic nitrogen deposition could cause large-scale decreases in bNpp that could push these forests beyond their adaptive capacity.

**Significance Statement:** Trees can allocate up to 80% of photosynthetically-produced carbon belowground, yet little is known about the environmental variables that control this allocation, limiting accurate assessment of how forests will respond to environmental change. We examined how climatic and soil variables affected belowground productivity and allocation, and found that soil dynamics, not climate, were the primary drivers of belowground carbon dynamics. Within biomes, environmental drivers of belowground allocation varied for forests with low versus high belowground allocation, indicating that environmental drivers are site-specific and the development of a within-biome classification system could be useful. Soil variables caused large shifts in primary productivity and allocation in boreal, but not cold temperate biomes, suggesting that boreal forests may exhibit lower resilience to environmental change.

## Introduction

Forest ecosystems in the Anthropocene face multiple and compounding stressors, including changes in climate and soil chemistry, shifts in natural disturbance regimes, and anthropogenic activities such as deforestation (Case et al. 2021, Johnson and Johnson et al. 2019, Lilleskov et al. 2024, Popkin 2019). Forests have adapted to their ecological and geographical environmental niches through within-plant carbon (C) allocation shifts or shifts in their landscape distribution (Körner 2015, White et al. 2016, Pagel et al. 2020). However, in the current climate crisis, many ecosystems are being pushed towards their biological tipping point as plants struggle to adapt to modified growing conditions (Pereira and Viola 2018, Hoegh-Guldberg et al. 2019). To assess how well a forest adapts to different stressors, there is a need for a more comprehensive understanding regarding the current productive capacity of forests, their genotypic and phenotypic adaptation to changing soil and climatic environments, and ecosystem fit – a measure of how well a forest is adapted to its site, defined as the ratio of field-collected total net primary productivity (tNpp) to the theoretical maximum tNpp (Gordon et al. 1983, Gordon et al. 1992).

Furthermore, it is important to understand how changes in tree C allocation to the belowground components impact the acquisition of growth-limiting resources (Vogt et al. 1983, Clark et al. 2001, Scurlock et al. 2002). However, accurate estimates of a forest’s current annual productive capacity, or net primary productivity (Npp), have been difficult to obtain (Clark et al. 2001, Scurlock et al. 2002). This has been due to poor predictions of belowground Npp (bNpp), which constitutes a significant portion of the total productivity of a forest and is therefore important to include in models estimating ecosystem C budgets. Yet, bNpp still remains the least known C pool in the C cycle (Vogt et al. 1993, Gherardi and Sala 2020, Klock et al. 2022a). This knowledge would provide further insights into how edaphic (factors relating to the structure and composition of soil) and climatic growth environments determine how much C is allocated to belowground tree components and whether restoring a site’s primary productive capacity is realistic (Klock et al. 2022a).

Although studies focusing on bNpp and edaphic and climatic conditions exist (Xiao et al. 2023), the results are challenging to use in building a holistic model to explore the sensitivity of bNpp to edaphic and climatic conditions (Vogt et al. 2016, Klock et al. 2022a). Individual field studies across biomes are inconsistent and have found opposing effects of environmental variables on bNpp (Gherardi and Sala 2020), and therefore cannot be compared to develop general relationships of factors impacting bNpp since they do not include a comprehensive suite of both climate and edaphic variables (but see Vogt et al. 2016). This suggests that any relationships found may not be generalizable or realistically represent the diversity of forests found in a biome. For example, in boreal forests, bNpp is typically limited by both belowground and aboveground resources due to temperature-limited nutrient mineralization rates, such as belowground nutrient availability or solar radiation (Gherardi and Sala 2020). In contrast, in a temperate deciduous forest, soil moisture and nutrient availability determined bNpp levels (Newman et al. 2006). In tropical forests, bNpp is affected by limiting aboveground factors such as light and temperature (Gill and Finzi 2016, Nemani et al. 2003). However, Klock et al. (2022a) showed that tropical forests had significantly lower % allocation to bNpp in the Paleotropics compared to the Neotropics due to the more nutrient-poor soils of the Neotropics. In addition, a study assessing forest research sites across the tropics showed that the relationship between precipitation and total Npp (tNpp) varied for three soil textural classes, likely due to the differing soil moisture retention capabilities of these soils which then affect soil organic matter and nutrient availabilities. These results support the potential development of relationships between tNpp and certain soil texture classes, but not for all tropical forests. Finally, most current Earth system and productivity models assume climate is the primary driver of productivity and C dynamics, even though studies increasingly show that soil plays an important role in these dynamics (Luo et al. 2019, Todd-Brown et al. 2013).

Prior to 2000, bNpp estimates were based on field measures of changes in root biomass over time, but many indirect measures (*e.g.*, ecosystem models and proximate variables) emerged after this time because of the need to include bNpp estimates for many sites that did not have prior root research. A common indirect approach to model C allocation to the belowground was based on aboveground measures, such as aNpp:bNpp, which did not effectively capture how plants adapt to their site edaphic and climatic conditions at the plot scale (Klock et al. 2022a).

Research has shown that C allocated belowground frequently cycles at different rates and is governed by different drivers than aboveground C, therefore these indirect estimates may generalize changes in forest growth at larger scales but would not adequately represent the diversity of site scale forest adaptations to environmental changes (Fortier et al. 2019, Klock et al. 2022a). Also, researchers lacking site-specific robust estimates of bNpp use indirect measures to calculate this variable based on relationships developed by other studies. The estimates of bNpp produced from direct and indirect methods often differ significantly, and it remains challenging to use indirect methods to identify the upper bounds of C available for root and mycorrhizal growth due to insufficient data availability from representative sites. To identify the mechanisms driving belowground productivity and allocation, we need to quantify the effect of a larger and more nuanced suite of edaphic and climatic variables on field-measured estimates of bNpp. This will allow for more accurate predictions of tNpp and a more detailed, quantifiable understanding of the biome scale patterns and drivers of bNpp and the % allocated to bNpp.

To better understand and identify the mechanisms controlling bNpp – the least understood part of the global C budget – we used random forest regression models to predict measures of tNpp, aboveground Npp (aNpp), bNpp, % allocation to bNpp, and ecosystem fit at two eco-physiological scales: phenological traits (evergreen, deciduous, mixed) and regional climatic zones (boreal, cold temperate), using a suite of 21 climate and edaphic variables in forests (Vogt et al. 1986, Vogt et al. 1995, Vogt et al. 1996, Klock et al. 2022a) and field-collected productivity measures. We hypothesized that climatic factors would primarily limit boreal forest tNpp, since tree growth is generally constrained by duration of the growing season, while temperate forests would be more limited by soil nutrients since a milder climate has less impact on forest productive capacity. To understand the drivers of bNpp, allocation, and ecosystem fit, our study main objectives were to:

i. determine how bNpp, % allocation to bNpp, and ecosystem fit vary with biome and phenology,
ii. assess the relative importance of climate vs. edaphic variables as predictors of bNpp, % allocation to bNpp, and ecosystem fit, and
iii. evaluate the response of bNpp, % allocation to bNpp, and ecosystem fit to soil and climatic drivers, and how this response varies between boreal and temperate biomes.

## Result and Discussion

### What are the patterns of bNpp and % allocation to bNpp in cold temperate and boreal biomes?

We determined that bNpp differed by biome and phenology. Specifically, boreal evergreen forests had significantly lower bNpp than boreal deciduous forests and cold temperate evergreen forests (Figure S1; Figure S2). This difference could be due to the high nutrient retention of boreal evergreen species because of their long leaf longevity (up to 10 years for *Picea glauca* and *Picea mariana;* Reich et al. 2014), which would reduce the % allocation to bNpp, as annual nutrient requirements are lower than for boreal deciduous species, or cold temperate evergreens. In contrast, percent allocation to bNpp did not vary by phenology or biome, or when examining phenology by biome or biome within phenology (Table 1; Figure S1; Figure S2). Previous studies have shown variable and overlapping values for % allocation to bNpp between boreal and temperate forests (Gower et al. 2001, Raich and Nadelhoffer 1989).

**Table 1.**
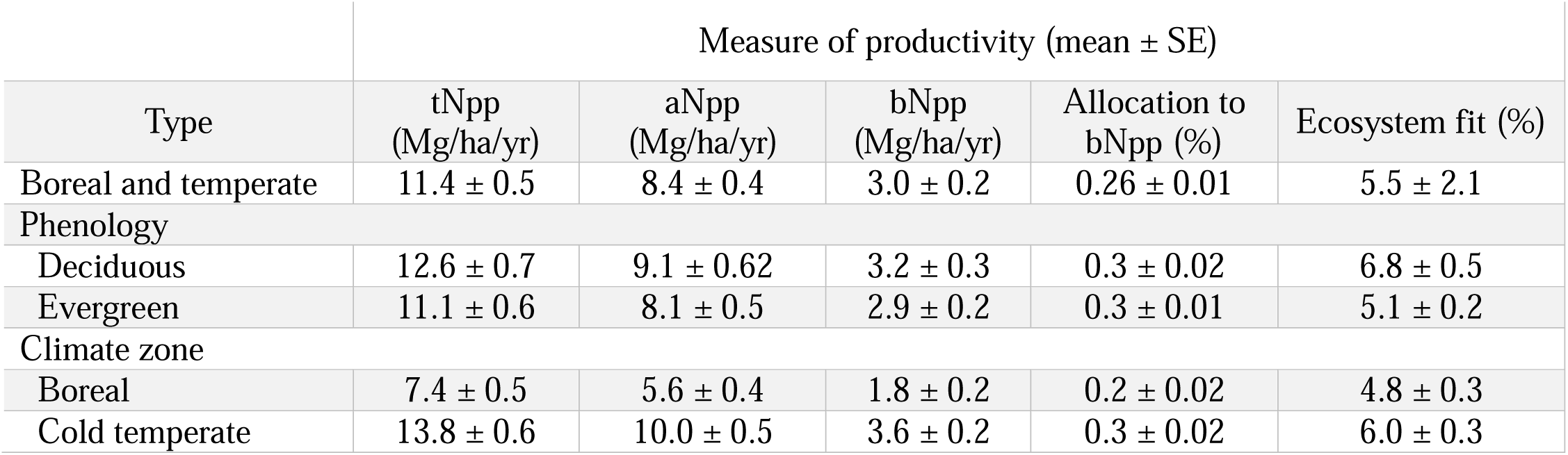
Summary statistics of Aboveground Net Primary Productivity (aNpp) and Belowground Net Primary Productivity (bNpp), % allocation belowground (% allocation to bNpp), and ecosystem fit.

| Type | Measure of productivity (mean ± SE) | | | | |
| --- | --- | --- | --- | --- | --- |
|  | tNpp<br>(Mg/ha/yr) | aNpp<br>(Mg/ha/yr) | bNpp<br>(Mg/ha/yr) | Allocation to<br>bNpp (%) | Ecosystem fit (%) |
| Boreal and temperate | 11.4 ± 0.5 | 8.4 ± 0.4 | 3.0 ± 0.2 | 0.26 ± 0.01 | 5.5 ± 2.1 |
| Phenology |  |  |  |  |  |
| Deciduous | 12.6 ± 0.7 | 9.1 ± 0.62 | 3.2 ± 0.3 | 0.3 ± 0.02 | 6.8 ± 0.5 |
| Evergreen | 11.1 ± 0.6 | 8.1 ± 0.5 | 2.9 ± 0.2 | 0.3 ± 0.01 | 5.1 ± 0.2 |
| Climate zone |  |  |  |  |  |
| Boreal | 7.4 ± 0.5 | 5.6 ± 0.4 | 1.8 ± 0.2 | 0.2 ± 0.02 | 4.8 ± 0.3 |
| Cold temperate | 13.8 ± 0.6 | 10.0 ± 0.5 | 3.6 ± 0.2 | 0.3 ± 0.02 | 6.0 ± 0.3 |

Generally, % allocation to bNpp increases with latitude from tropical to temperate to boreal forests, due to temperature-limited mineralization in higher latitudes that constrains belowground resources and induces greater allocation belowground to acquire limiting nutrients needed for growth and maintenance (Gherardi and Sala 2019). Boreal forests are thus typically more nutrient-limited than temperate forests due to slower rates of decomposition and nutrient cycling. Therefore, they are usually more dependent on symbiotic associations (*e.g.*, mycorrhizal fungi) to acquire growth limiting nutrients (Anthony et al. 2022), which leads to higher rates of belowground C allocation to roots and mycorrhizas (Vogt et al. 1982). This pattern, however, was not supported by a larger database where boreal forests had a significantly lower belowground biomass than temperate or tropical biomes (Klock et al. 2022a). Other studies have found that temperature-limited mineralization drives belowground resource limitation in both boreal and temperate forests in similar ways (Kicklighter et al. 2019). These differences in belowground allocation between studies show that biome-scale patterns of % allocation to bNpp are not consistent and suggest that many belowground studies do not adequately describe the role of mycorrhizas and C allocation to fungal biomass to acquire nutrients. However, Clemmensen et al. (2013) reported that roots and root-associated microorganisms are a dominant source of soil organic matter in boreal forests. Additionally, the functional balance theory proposes that plants allocate their resources to minimize resource limitation to growth and therefore gain access to further resources (Gherardi and Sala 2019). Therefore, plants generally increase the fraction of productivity allocated belowground with decreasing resource availability, such as drought conditions or soil nutrient limitation. The lack of difference in the % allocation to bNpp between boreal and temperate forests suggests that belowground resources are limiting in a way that similarly affects allocation patterns in both forests. However, it could also indicate that local site-scale factors and local site variability are primary drivers of % allocation to bNpp, and that biome-scale patterns are not sensitive to variation in these environmental variables, as shown by Klock et al. (2022a).

A wide variety of environmental conditions may influence allocation patterns, and site-specific factors can be more important than biome-scale differences when comparing C allocation patterns across forest types (Gower et al. 2001). Therefore, some authors caution against making broad generalizations based on forest type alone, which is further supported by the wide range of values reported in estimates of % allocation to bNpp (Körner 2003). For example, some studies estimate that % allocation to bNpp in boreal forests ranges from 45-65%, while others report 25-30%, whereas temperate forests generally fall within the range of 30-46% (Gill and Finzi 2016, Gower et al. 2001, Raich and Nadelhoffer 1989). Similar reasoning can be applied to the similarity we found in % allocation to bNpp between deciduous and evergreen forests in this study. Although some studies show that evergreen forests allocate a higher fraction of Npp belowground than deciduous forests (Gherardi and Sala 2020, Gower et al. 2001), our study shows that there can be considerable site-scale variation in % allocation to bNpp, particularly for evergreen forests (Figure S1).

When examining the distribution of % allocation to bNpp in boreal and temperate zones, we found that boreal % allocation to bNpp exhibited a bi-modal distribution, with clear peaks at 15% and 45%, suggesting site-scale variability and clustering in % allocation to bNpp within the boreal climate type. Cold temperate forests, on the other hand, had a mean % allocation to bNpp of 27%; however, the range for % allocation to bNpp was 20% higher for cold temperate (5-83%) than boreal (5-62%) forests, with some evergreen cold temperate forests allocating 83% of tNpp belowground (Table 1, Figure S1). The lack of difference between the two biomes and the larger range of % allocation to bNpp suggests that cold temperate forests have higher variability in their growing environmental conditions and site variation than boreal forests. This also further confirms that within-biome site-scale variation is important to consider when evaluating the drivers of belowground allocation patterns.

The very high values of % allocation to bNpp (60-80%, Figure S1; Figure S2) in cold temperate forests suggest release from aboveground resource limitation such as precipitation, leading to higher % allocation to bNpp due to belowground nutrient limitation to growth. Previous evidence suggests that bNpp in boreal forest systems is not driven by precipitation but controlled by nutrient limitation or solar radiation, indicating colimitation of above- and belowground resources to growth (Gherardi and Sala 2019). Tropical forests are also influenced by both above and belowground resource limitation, and especially by the diversity of growth limiting microsites (Klock et al. 2022a). Temperate forests, on the other hand, tend toward the average of tropical and boreal systems in productivity and diversity (Klock et al. 2022a, Reich and Bolstad 2001) and have milder climates and more variability in climate and geography. Thus, they are limited more by belowground resources as they are released from the dominance of aboveground resource limitation.

### How does ecosystem fit vary with forest biome and phenology?

Ecosystem fit was significantly higher for deciduous forests than evergreen forests. Ecosystem fit was also significantly higher in temperate than boreal forests, but only for evergreen forest types (Table 1, Figure S1). Ecosystem fit is the ratio of field-collected tNpp to the theoretical maximum tNpp and estimates how well a forest is adapted to its site and whether it has the capacity to grow more biomass beyond its current growth rate indicated by direct field measures of tNpp (Gordon et al. 1983, Gordon et al. 1992). Ecosystem fit depends on solar radiation and photosynthetic efficiency and is independent of local soil conditions, thus providing an independent measure of a forest’s productive capacity (Klock et al. 2022b). Therefore, it is not surprising that deciduous traits typically involve higher photosynthetic efficiency, as the amount of total solar radiation generally limits boreal forest productivity. Importantly, phenology mediated the effect of forest biome on ecosystem fit. The overall ecosystem fit of deciduous trees was higher and did not differ between forest biomes, whereas the ecosystem fit of evergreen trees was lower and was influenced by forest biomes (Table 1, Figure S1), suggesting that deciduous forest types are more resilient to climate influences whereas evergreen traits are more sensitive to large-scale variations in climate.

### What is the relative importance of soil variables versus climate variables in predicting belowground productivity, allocation, and ecosystem fit?

Soil variables, particularly soil nutrient flux rates, were the most important variables (Figure 1, Table S1). The effect of climate was secondary to the effect of soil variables (specifically soil flux rates), within and across biomes (Table 2). This suggests that soil characteristics, rather than climate variables, are the most important predictors of belowground productivity, allocation, and ecosystem fit. These results differ from a related study by Klock et al. (2022a), which found that a combination of climatic and edaphic variables explained productivity. However, Klock et al. (2022a) used a limited number of edaphic variables consisting of soil texture and soil taxonomic order, whereas this study employed a broad suite of aboveground and belowground soil variables which provided a more detailed analysis of specific environmental drivers of belowground productivity. The top five most important variables consisted of four soil flux rates (aboveground litter phosphorus (P) flux, soil nitrogen (N) flux, forest floor P mean residence time, and aboveground litter N flux), and one soil pool (forest floor P). The most important variables across and within biomes were P and N flux rates, suggesting colimitation of both N and P on productivity.

**Figure 1.**
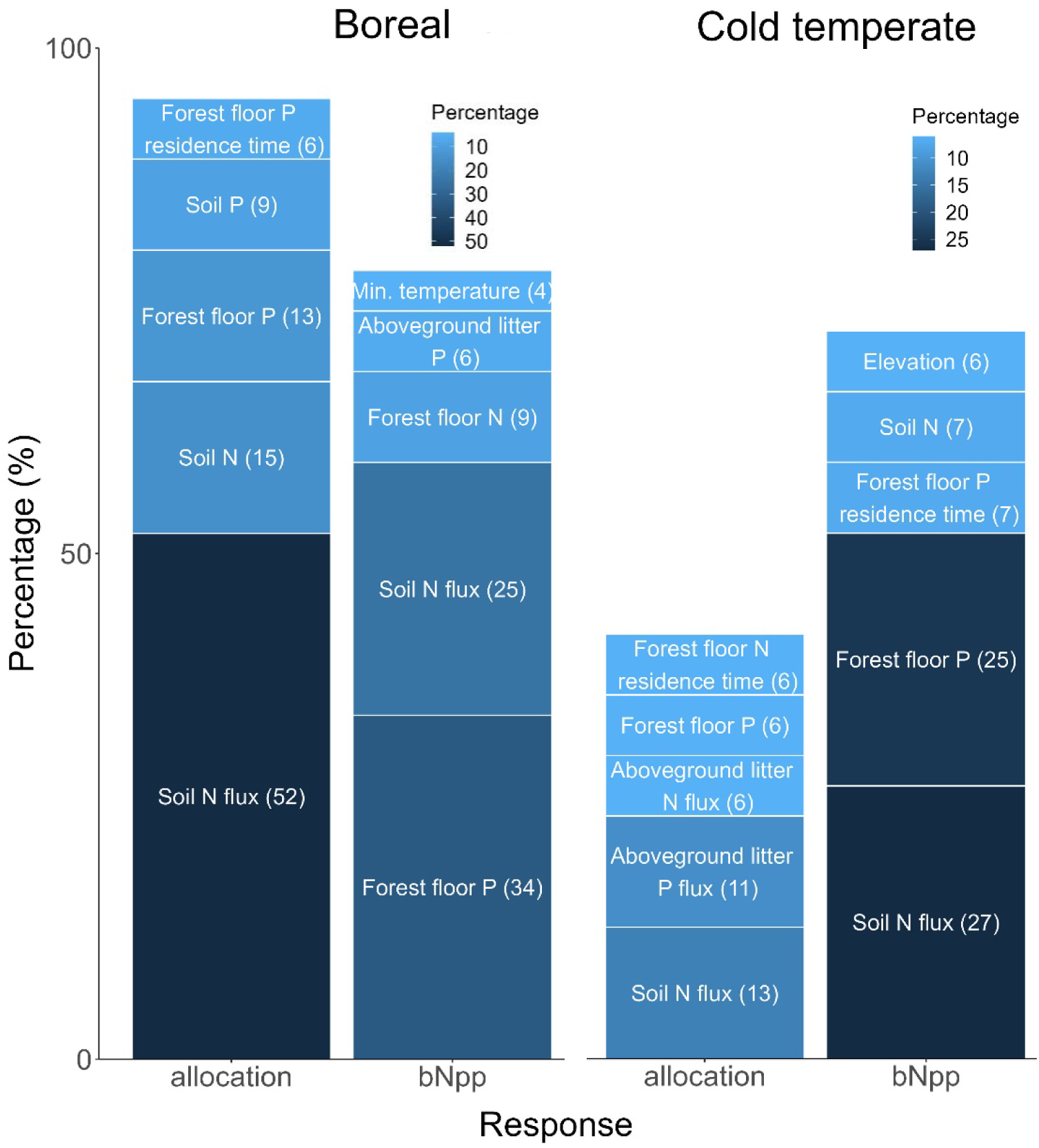
The top five most important variables identified by random forest regression models predicting belowground net primary productivity (bNpp) and % allocation to bNpp, for boreal and cold temperate forests. For each model, the top five variables are stacked in order of importance. The height of each stack within the boxplot corresponds to the percentage of the variable’s importance for each model, and % variable importance is written alongside each variable.

**Table 2.** The % relative importance of predictor variables grouped by climate (mean air temperature, minimum air temperature, maximum air temperature, total annual precipitation), soil properties (soil taxonomic order, soil texture), soil pools (soil N, soil P, soil organic matter, forest floor mass, forest floor N, forest floor P), and soil fluxes (aboveground litter flux, belowground litter flux, aboveground litter N flux, aboveground litter P flux, soil N flux, forest floor mean residence time, forest floor N mean residence time, forest floor P mean residence time) for random forest models estimating the effects of these variables on the response variables of tNpp, bNpp, % allocation to bNpp, and ecosystem fit in boreal and cold temperate forests.

|  | Predictor variable groups (% relative importance) |  |  |  |  |
| --- | --- | --- | --- | --- | --- |
|  | Response | Climate | Soil properties | Soil pools | Soil fluxes |
| Boreal | tNpp | 14 | 3 | 25 | 58 |
|  | bNpp | 14 | 3 | 51 | 33 |
|  | % Allocation to bNpp | 4 | 1 | 22 | 72 |
|  | Ecosystem fit | 20 | 10 | 22 | 47 |
| Cold temperate | tNpp | 20 | 4 | 23 | 53 |
|  | bNpp | 15 | 5 | 43 | 38 |
|  | % Allocation to bNpp | 23 | 6 | 27 | 45 |
|  | Ecosystem fit | 19 | 2 | 11 | 67 |

The effects of soil flux variables on productivity measures indicated that aNpp, tNpp and ecosystem fit increased with soil flux rates and soil pool size (Table S2). This is an expected pattern as productivity generally increases with nutrient cycling rates and availability as nutrient limitation is released and fuels growth (Wieder et al. 2015). However, these trends did not hold for bNpp and % allocation to bNpp. Individual soil flux rates and soil pools had differing and sometimes opposite effects for bNpp and % allocation to bNpp, and between measures of below- and aboveground productivity (Figure 2, Table S2). The fact that bNpp and % allocation to bNpp do not follow the same patterns as aboveground- or total productivity indicates that belowground processes are governed by different drivers than aboveground processes (Fortier et al. 2019), further emphasizing the need to fully understand and unravel the interactions between belowground processes, especially those of fungal symbionts, which are an important driver of soil nutrient cycling (Averill et al. 2014).

**Figure 2.**
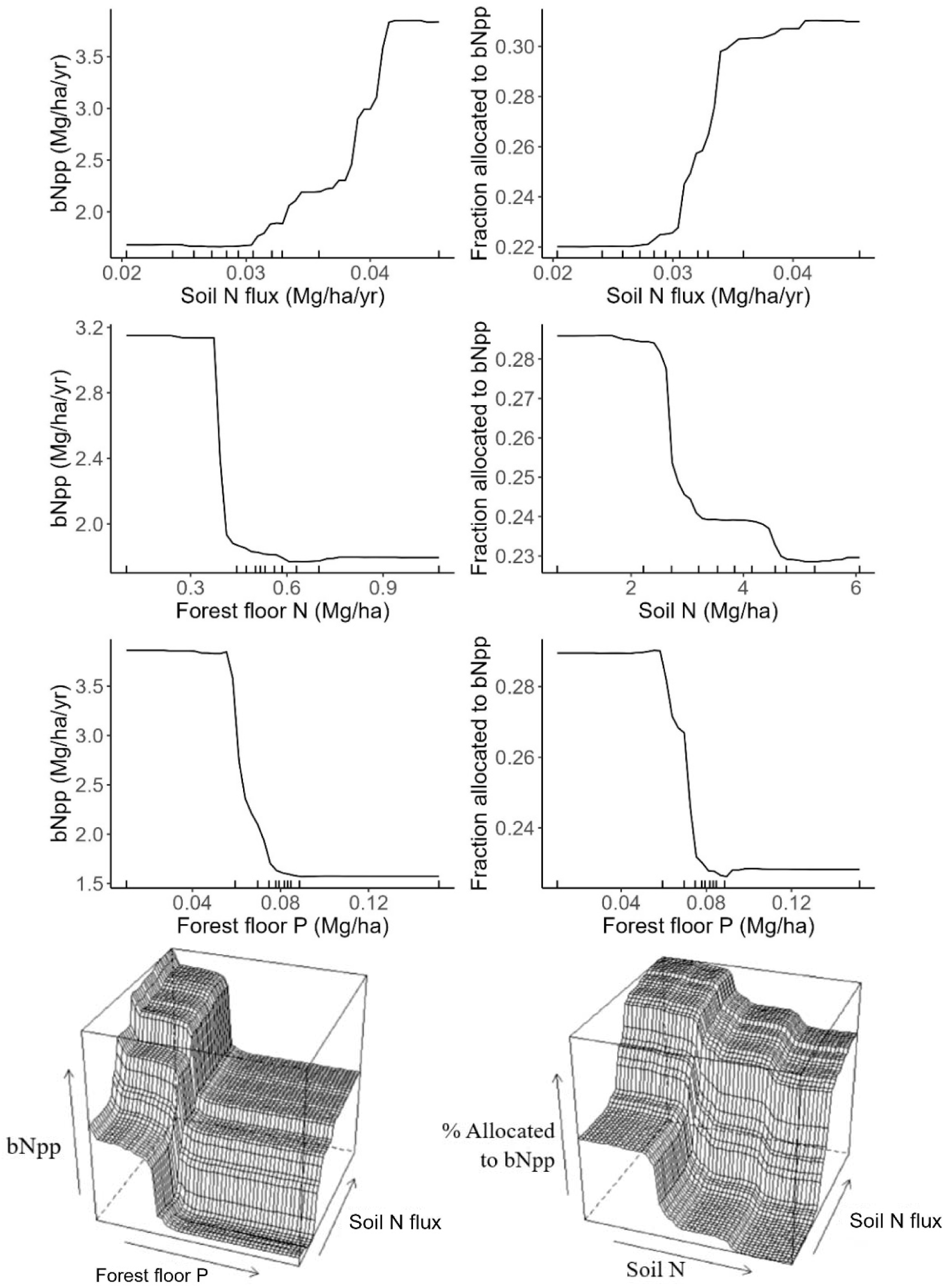
Partial dependence plots for boreal forests. Left panels illustrate the marginal effects of the most important variables on belowground net primary productivity (bNpp), which were soil N flux (top panel), forest floor N (second panel) and forest floor P (third panel). Right panels illustrate the effect of the most important variables on % allocation to bNpp, which were soil N flux (top panel), soil N (second panel), and forest floor P (third panel). The bottom panels show a three-dimensional partial dependence plot illustrating the effect of forest floor P and soil N flux on bNpp, and the effect of soil N and soil N flux on % allocation to bNpp. Tick marks represent data deciles.

### How did soil variables affect boreal forest productivity, allocation, and fit?

Soil flux variables had especially high variable importance for boreal forests, particularly for allocation belowground, where soil flux variables accounted for 72% of the relative importance of all environmental variables (Table 2). This was driven primarily by soil N flux and the soil N pool, which constituted 60% of the relative importance of all variables (Figure 1), indicating that these are main drivers of allocation belowground in boreal forests. Belowground productivity was primarily constrained by soil N flux and the forest floor P pool, also accounting for 60% of the relative variable importance (Figure 1). Allocation to the belowground was thus primarily limited by N, while bNpp was co-limited by N and P. Recent studies have provided new evidence that belowground productivity in boreal forests is not controlled by precipitation but driven by nutrient limitation or solar radiation (Gill and Finzi 2016, Nemani et al. 2003), supporting our results.

Our results also indicate that a few drivers constrain the system in unique ways. For example, the rate of soil N flux increased allocation belowground dramatically, with a hard threshold effect (sudden and large change) at 0.031 Mg/ha/yr that drove an increase in belowground allocation from 22% to 31% (Figure 2, Figure 1, Table S3). This indicates that soil N flux had a narrow range of effect on allocation but that effect on % allocation to bNpp was relatively strong. Furthermore, this suggests the existence of two states of belowground allocation in boreal forests – a low and high allocation state, driven by a critical environmental threshold determined by soil N flux rate. The effect of the soil N pool on allocation seems to confirm the presence of two distinct states of % allocation to bNpp – allocation decreases at a hard threshold of 2.6 Mg/ha of soil N, with a range of effect on % allocation to bNpp from 29% to 23% (Figure 2, Table S3). This also suggests that boreal forest allocation is split between two systems – one with low allocation (characterized by low rates of soil N cycling and a high soil N pool), and one with high allocation (characterized by high rates of soil N cycling and a low soil N pool). The high soil N flux at sites with high belowground allocation could reflect increased fungal activity, where low soil N levels would demand increased C allocation from tNpp (Wallenstein et al. 2006). The bi-modal distribution of % allocation to bNpp with clear peaks at 15% and 45% further confirmed this, suggesting site-scale variations and clustering in % allocation to bNpp within the boreal climate type.

For boreal bNpp, productivity increased with soil N flux and decreased with the forest floor N and P pools, and although hard threshold effects of a few dominant soil variables drove changes in bNpp, these changes were less impactful than those for % allocation to bNpp (Figure 2). The presence of hard threshold effects and the high % variable importance of just two soil variables suggests that belowground productivity and allocation in boreal forests have low adaptive capacity or resiliency (Seddon et al. 2011) and lack the ability to adapt to different environmental conditions within the variability of site variables. Furthermore, this suggests that these forest types could be prone to sudden changes in bNpp and % allocation to bNpp due to critical thresholds in soil flux rates, confirmed by the significantly lower ecosystem fit of boreal forests compared to temperate forests (p < 0.001; Figure S1).

### Are soil variables driving % allocation to bNpp in boreal forests site-specific?

To further understand the drivers of these within-biome, site-scale groupings, we used random forest regression models to examine the effect of climate and edaphic variables on boreal forest systems characterized by low (≤ 20%) and high (> 20%) allocation patterns, clustered by the “sidClustering” function within the randomForestSRC package in R (Mantero and Ishwaran 2021, R Core Team 2020; Table S4). Results showed that boreal forests characterized by low belowground allocation were driven primarily by N-associated edaphic variables, while boreal forests with high belowground allocation were controlled by P-associated variables (Table S4; Figure S3). This suggests that belowground allocation in boreal forests is both N and P limited, but site-scale conditions drive these limitations and these differences were only observable within biomes. Furthermore, when examining the effects of dominant environmental drivers (soil N pool and soil N flux) for forests with high vs. low belowground allocation, we found that abrupt and large decreases in allocation were found in boreal forest sites with low allocation, and not in sites with high allocation (Figure S4). These abrupt allocation changes indicate that boreal forest stands with low belowground allocation are especially sensitive to changes in the environment and prone to sudden shifts in allocation with changing edaphic factors. For boreal forest productivity, the primary environmental drivers for low productivity sites (<2 Mg/ha/yr Npp) were aboveground litter N flux and aboveground litter flux, while for high productivity sites (> 2 Mg/ha/yr Npp) the dominant drivers were forest floor P and the soil P pool (Figure S5). These results demonstrate that scale is an important consideration in understanding drivers of productivity necessary for local forest assessments and management, especially because recent studies have confirmed that belowground allocation is highly variable within biomes, resulting in high uncertainty in global estimates of belowground processes. Furthermore, site-specific drivers of % allocation to bNpp provides evidence that plant communities and C allocation patterns adapt to local environmental site conditions (Luo et al. 2019).

### How did soil variables affect cold temperate forest productivity, allocation, and fit?

Belowground productivity and allocation in cold temperate forests were driven primarily by soil pools and fluxes, although climate had a higher relative importance overall for productivity in cold temperate compared to boreal forests (Table 2). The fraction of productivity allocated belowground was driven by multiple soil variables, dominated primarily by soil flux rates, with relatively even variable importance (Figure 1). These variables were also split relatively evenly between aboveground and belowground N and P flux rates, suggesting colimitation of both N and P on allocation, which differed from boreal forests where N limitation seemed to be the main driver of % allocation to bNpp. Further confirming the differences in nutrient limitation, soil N was the dominant soil pool variable driving belowground allocation in boreal forests, while forest floor P was the dominant soil pool variable explaining allocation for cold temperate forests.

The large number of variables with similar percent values of relative importance for cold temperate forests suggests that there were no dominant drivers of % allocation to bNpp, with influence spread across many variables. Generally, soil variables affected allocation over a much wider range than in boreal forests, with more gradual non-threshold effects (Figure 1; Figure 3). However, soil variables had a much lower magnitude of effect on % allocation to bNpp in cold temperate forests (25-30%) than boreal forests (20-40%), indicating that allocation in boreal forests was much more sensitive to changes in soil nutrient cycling, and that these changes were more abrupt and intense.

**Figure 3.**
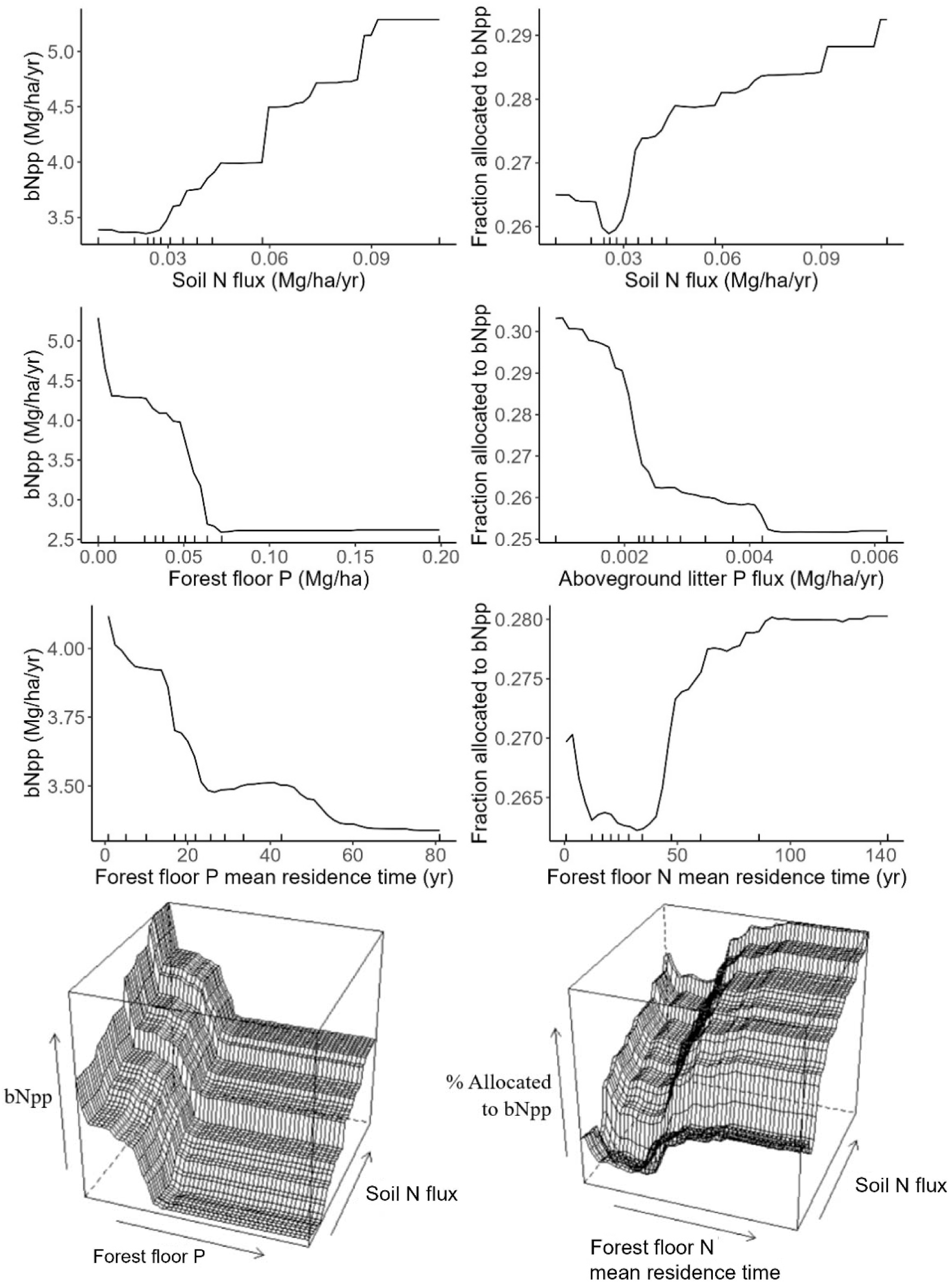
Partial dependence plots for cold temperate forests. Left panels illustrate the marginal effects of the most important variables on belowground net primary productivity (bNpp), which were soil N flux (top panel), forest floor P (second panel), and forest floor P mean residence time (third panel). Right panels illustrate the effect of the most important variables on % allocation to bNpp, which were soil N flux, aboveground litter P, and forest floor N mean residence time. The bottom panels show a three-dimensional partial dependence plot illustrating the effect of forest floor P and soil N flux on bNpp, and the effect of forest floor N mean residence time and soil N flux on % allocation to bNpp. Tick marks represent data deciles.

Some variables had gradual non-monotonic effects on % allocation to bNpp, where a change in the direction of effect on the response occurred. For example, soil N flux decreased % allocation to bNpp for lower values of its range, but that effect was reversed for higher values of soil N flux, with trend change occurring at 0.025 Mg/ha/yr (Figure 3). The same trend was present for forest floor N mean residence time, and forest floor mean residence time (Figure 3), with % allocation to bNpp increasing for low rates of forest floor N flux and decreasing with high rates of forest floor N flux, the reverse of the trend observed for soil N flux on allocation. This illustrates that aboveground and belowground soil N cycling variables could have differing effects on % allocation to bNpp. These non-monotonic effects also suggest the presence of environmental thresholds that cause shifts in ecosystem properties.

Although some threshold-like effects were observed in several soil variables such as belowground litter flux and aboveground litter P flux, these were much less dramatic than those for boreal forests and indicative of some limitation release. These results show that cold temperate forests had many influential soil variables that affected belowground productivity and allocation in complex and non-linear ways, and with a large variability in responses. However, there was also the presence of some threshold values that indicated sudden changes or trend reversal in % allocation to bNpp. These results suggest that cold temperate forests, like boreal forests, exhibit a type of environmental threshold characterized by trend reversal. But unlike boreal forests, belowground productivity and allocation in cold temperate forests changed more gradually in response to environmental variables, suggesting higher adaptive capacity. This is further supported by the large number of variables with even variable importance, the lack of sharp threshold effects, and the lower magnitude of effect of soil variables on measures of productivity, all suggesting that bNpp and % allocation to bNpp in cold temperate forests are more resilient to changes in soil nutrient cycling. In addition, effects of soil variables on productivity within cold temperate forests was more variable and complex than those for boreal forests, indicating that cold temperate forests had higher variability of soil effects on productivity and a larger suite of interactions, making them potentially more adaptable to changes in the environment than boreal forests.

Soil variables for both boreal and cold temperate forests exhibited saturating effects on bNpp and % allocation to bNpp, where an increase in the soil variable had no effect on the productivity response variable. This suggests the presence of multiple limiting variables, where a variable limits productivity in the range of its effect, and limitation is removed upon saturation, whereupon some other environmental variable then limits the response variable. There is accumulating evidence that multiple (i.e., two or more) nutrient variable limitation is relatively common in forest ecosystems (Elser et al. 2007, Harpole et al. 2011, Fay et al. 2015). This colimitation can be defined as simultaneous colimitation, where both nutrients are needed to induce an increase in productivity, or independent colimitation, where productivity increase can be induced by either nutrient separately (Harpole et al. 2007), demonstrated in this study. These results further support the complex and non-linear interactions that regulate productivity and allocation.

## Conclusion

Soil variables were the most important predictors of belowground productivity and allocation, whereas climate effects were tertiary after soil nutrient flux rates and soil nutrient pool variables. Therefore, soil flux rates and pools must be explicitly modeled to accurately predict feedback between climate, productivity, and allocation. This is contrary to most current Earth system and productivity models where climate is assumed to be the primary driver of productivity and C dynamics (Luo et al. 2019, Todd-Brown et al. 2013), indicating that more explicit and detailed representation of soil variables (e.g., the expansion of soil modules to include additional soil flux variables) needs to be incorporated to increase the accuracy of productivity and allocation predictions within forest ecosystems. Recent novel studies on global bNpp have shown that large-scale climate variables such as precipitation do not drive changes in bNpp or % allocation to bNpp in forest ecosystems (Gherardi and Sala 2020). Another recent study emphasized similar importance for global subsoil organic C turnover times, which were dominantly controlled by soil properties rather than climate at both local and global scales, providing similar insight on a previously unmapped albeit crucial belowground soil property (Luo et al. 2019).

Nitrogen and P soil pool and flux rates were the most important variables for predicting productivity and allocation in boreal and cold temperate forests, indicating the colimitation of both N and P across forest types. The type of limitation varied when forest belowground productivity was further subdivided into low and high productivity categories, indicating that within-biome site-level evaluation is important. Belowground productivity and allocation within boreal biomes were driven by few variables characterized by hard threshold effects resulting in sudden and large changes in productivity measures, signifying low adaptive capacity. Conversely, belowground productivity and allocation in cold temperate forests were influenced by many variables, a lack of hard threshold effects of environmental variables on productivity, and gradual changes in response variables, indicating a higher capacity to adapt to environmental change. This indicates that cold temperate forests might be more resilient to climate change while boreal forests could be more sensitive and could be more prone to sudden changes in productivity and allocation in response to changes in climate or other disturbances such as prescribed fire (Buonanduci et al. 2020, Harvey et al. 2023, Johnson and Johnson et al. 2019) or anthropogenic N deposition (Holtgrieve et al. 2011, Lilleskov et al. 2024). The lower resilience of boreal forests was further supported by lower ecosystem fit, and clustering of bNpp and % allocation to bNpp that showed that a higher percentage (70%) of boreal forest sites had lower bNpp than cold temperate sites (57%, Table S4). Importantly, increases in forest floor N and soil N led to state shifts and large decreases in belowground productivity and allocation in boreal forests, therefore N deposition has the potential to cause large decreases in bNpp and potentially push these forests beyond their adaptive capacity. Other disturbances such as drought could alter productivity by decreasing belowground cycling of N, leading to large and abrupt decreases in belowground productivity.

However, boreal forests generally have the highest fungal diversity of all forest types (Li et al. 2022) and studies have reported strong associations between fungi and plant hosts, where mycorrhizal fungi can receive up to 20% of plant host C (Hobbie 2006, Konvalinková et al. 2017). Mycorrhizal fungal associations might help boreal forests maintain resilience to stressors such as drought and N deposition due to the ability of mycorrhizal fungi to supply water and mine nutrients for plant hosts.

Due to the lack of field-collected measures of productivity in a diversity of forest ecosystems globally, vegetation models are currently the best way to estimate C sequestration within forests and the effects that anthropogenic impacts and climate change will have on these C stores (Case et al. 2021). Using new modeling techniques in machine learning allows a more nuanced and detailed understanding of the mechanisms underpinning the controls on forest productivity, allocation, and adaptive capacity, specifically modeling non-linear effects and identifying thresholds of productivity and how these are driven by environmental variables, which are not able to be captured by traditional ecosystem vegetation models. Machine learning models can provide crucial information to forest management, such as quantifiable thresholds for biome-scale productivity at which state changes occur. In addition, our results showed that environmental drivers differed for varying productivity levels within biomes, indicating that development of within-biome classifications for forest systems could prove useful for management activities. Estimates of ecosystem fit can also allow managers to better understand a forest’s productive capacity and predict whether forests can achieve higher levels of productivity and what drives the structure and function of these forests at the ecosystem scale. Although this information is important to assess and maintain forest health in response to climate change and other anthropogenic stressors, this research is also needed to inform communities that depend on forests to survive. Understanding the environmental drivers of a forest’s productivity, allocation and ecosystem fit will determine if these forest ecosystems can provide ecosystem services for populations that rely on these forests (Chao 2012), and if forests can adapt to climate change within the constraints of edaphic controls. Our results allow for a more fine-scale assessment of forest regions sensitive to climate change, and further help guide site-specific policies and management for forest ecosystem health and effective C management. Furthermore, these results are essential to understanding and modeling the C cycle, especially as bNpp is the least known component of global C dynamics.

## Materials and Methods

### Dataset and variable selection

We used a unique and extensive multiple-biome dataset of field-based measurements conducted over 40 years to test the questions posed for this research (Table S6, Klock et al. 2022a). The dataset contained soil and climate variables, and tNpp, aNpp, and bNpp data collected per site (collection methodology is described in the literature listed in Table S6). The dataset described 181 natural forest sites, which were characterized at two spatio-ecological scales: leaf phenology types at the large scale (139 evergreen; 42 deciduous) and climatic regions at the meso-scale (69 boreal – 84% evergreen; 99 cold temperate – 74% evergreen). Climatic and soil variables are listed in Table S1. By phenology and regional climate groupings, almost all soil and climate variables were used to predict ecosystem fit, tNpp, aNpp, bNpp, and % allocation to bNpp. The soil variables aboveground litter flux and belowground litter flux were only used to predict ecosystem fit, as these variables include production measures. A previous study examined this dataset with a focus on tNpp and ecosystem fit and used an expanded dataset that contained additional biomes such as tropical forests (Klock et al. 2022a). Our study provides a detailed look at sites that contain a larger suite of soil variables to comprehensively assess drivers of belowground productivity, allocation, and ecosystem fit.

We used ecosystem fit as a response variable to assess how well a forest was adapted to its site constraints to grow biomass. Ecosystem fit uses a reference maximum productivity level calculated with factors external to the site and not influenced by internal site conditions (Gordon et al. 1992, Gordon et al. 1995). The most important climatic and edaphic variables can be used to identify the “safe operating space” at each site; that is, where the upper and lower bounds of environmental conditions will determine the forests that are better adapted to their current growth environment and which forests might be more vulnerable to climate change and may need management intervention (Johnstone et al. 2016, Scheffer et al. 2015). Ecosystem fit uses the Theoretical Maximum Productive Capacity, which is modeled as the product of mean solar radiation (Fick and Hijmans 2017), interception efficiency, the efficiency with which forest canopies absorb solar radiation, and the conversion efficiency, or the rate at which solar radiation is absorbed by plants and converted to biomass (Delucia et al. 2014). Model parametrization, equations, and values were taken from Klock et al. (2022b). This model assumes that photorespiration is a constant fraction of photosynthesis, photosynthetically active radiation is 45% of total solar radiation, forest canopies absorb 90% of incoming photosynthetically active radiation during active growth, and growth is not limited by water or nutrients.

### Model comparison and selection

We conducted model comparisons between linear regression (R package “MASS”), conditional forest (R package “party”), and random forest (R package “randomForest”) (R Core Team 2020) models using all soil and climatic variables to predict tNpp (Table S5). Although linear regression models are easier to interpret, random forest models work well with large datasets, work with missing data, and work better with non-linear trends (Lemon et al. 2003), which we identified with these data. Random forest models showed the lowest root mean squared error and mean absolute error for all measures of productivity (tNpp, aNpp, bNpp, % allocation to bNpp, and ecosystem fit). The random forest regression models were then used to predict the effect of soil and climatic variables on aNpp, bNpp, % allocation to bNpp, and ecosystem fit for boreal and cold temperate forests. For each model, we calculated root mean squared error (RMSE), % variance explained by the predictor variables (pseudo R^2^), the mean squared error of out-of-bag errors, and model mean absolute error (MAE) (Table S2), as well as the percent relative importance of climatic and soil variables (Table 2) and the most important variables based on %IncMSE (Figure 1). To evaluate the overall relative importance of soil and climate variables, we summed the relative importance of individual variables for soil and climate, respectively.

This study used the “randomForest” package in the R-environment (Liaw and Wiener 2002, R Core Team 2020). The algorithm combines large sets of decision trees formed by selecting sets of variables to improve classification and regression analysis. The recursive feature elimination function was used to identify the most important predictor variables with respect to the response variable. For modeling, 2/3 of the samples were used for training the algorithm, and 1/3 (‘out of bag’ or OOB samples) were used for cross-validation to determine the model error (OOB error). The major parameters required for proper optimization of the random forest model included: ‘ntree’, the total number of regression trees grown from bootstrap samples of the observations, ‘mtry’ – the number of predictor variables examined at each node, ‘nodesize’ – the smallest size of the end nodes of the trees grown. Following multiple iterations, the optimum ‘ntree’ selected was 500 at ‘mtry’ = 8 as it resulted in the smaller OOB error. To avoid overfitting, we optimized a tuning parameter that governs the number of features that are randomly chosen to grow each tree from the bootstrapped data using K-fold cross-validation, where K ∈ {5,10}, and chose the tuning parameter that minimizes test sample prediction error. In addition, growing larger forests improved predictive accuracy, although there are usually diminishing returns once there are up to several hundred trees. Further information regarding partial dependence plots, multicollinearity and unbiased variable selection can be found in the SI.

## Acknowledgments

We acknowledge the work of all researchers that collected these data over the years. We acknowledge Dr. Erik Lilleskov for reviewing this manuscript. AMB was supported by the H. Mason Keeler Endowed Professorship in Sports Fisheries Management.

## Author Contributions

Conceptualization, K.V. and D.V.; Methodology, K.V., D.V., A.K. and A.Y.P.; Statistical Analysis, A.K. and A.Y.P.; Investigation, K.V., D.V., A.K., K.M., A.M.B., and A.Y.P.; Writing & Editing, K.V., D.V., A.K., K.M., A.M.B., and A.Y.P.

