## Supplementary Information for "Forest belowground productivity and carbon allocation predominantly driven by soil properties rather than climate"

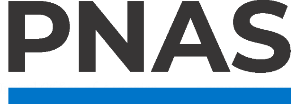

**Supporting Information for**

**Forest belowground productivity and carbon allocation predominantly driven by soil properties rather than climate**

Anne Y. Polyakov^1,2*^, Angela M. Klock^3^, Korena K. Mafune^4^, Andrew M. Berdahl^1,2^, Kristiina A. Vogt^3^, and Daniel J. Vogt^3^.

^1^Quantitative Ecology and Resource Management, University of Washington; Seattle, WA, 98195, USA.

^2^School of Aquatic and Fishery Sciences, University of Washington; Seattle, Washington, 98195, USA.

^3^School of Environmental and Forest Sciences, University of Washington; Seattle, Washington, 98195, USA.

^4^Department of Civil and Environmental Engineering, University of Washington; Seattle, Washington, 98195, USA.

**This PDF file includes:**

Supporting text

Figures S1 to S5

Tables S1 to S6

SI References

**Supporting Information Text**

**Model comparison and selection.** Results for random forest models typically consist of variable importance plots, and 1D and 2D partial dependency plots (Cutler et al. 2007). Variable (or feature) importance is the standard interpretation of random forest models and identifies variables with the best predictive power. The higher the importance, the more relevant the variable is according to the model. The mean decrease in accuracy, or percent increase in mean squared error, is the most robust and informative measure for the variable importance plot and shows how much model accuracy (model MSE) decreases if that variable is left out (or how much model accuracy increases if that variable is included). In other words, it is the increase in model MSE of predictions (estimated with out-of-bag-samples) as a result of variable j being permuted (values randomly shuffled). Partial dependency plots show the marginal effect that one (1D) or two (2D) features have on the predicted outcome of a machine learning model. A partial dependency plot can show whether the relationship between the target and a feature is linear, monotonic or more complex.

Random forest regression models have several advantages compared to linear models, since they do not require the typical assumptions of linear models to be met such as linearity, normality, no multicollinearity and homoscedastic variance. In addition, random forest models can work with large, unbalanced datasets with varying data types, are able to assess complex, non-linear interactions between variables and calculate variable importance (Lemon et al. 2003). Concerning correlated predictors in random forest models, multicollinearity is not an issue for random forest models. This is because each tree node is constructed by finding a single predictor and cut point, so only one candidate-predictor is examined at once. That is why relationships between predictors do not create problems since the random forest model never looks at more than one predictor at once. In addition, at each node, only a subset of predictors is taken into account, which is another anti-collinearity feature of random forest models.

However, although multicollinearity is not a problem for the random forest algorithm, it may be a problem for the random forest user. When the dataset has two (or more) correlated features, then from the model's point of view, any of these correlated features can be used as the predictor, with no concrete preference of one over the others. However, once one of them is used, the importance of others is significantly reduced, since the impurity they can remove is effectively already removed by the first feature. Consequently, they will have a lower reported importance. This is not an issue when we want to use feature selection to reduce overfitting, since removing features that are mostly duplicated by other features makes sense. But when interpreting the data, it can lead to the incorrect conclusion that one of the variables is a strong predictor while the others in the same group are unimportant, while they are actually very close in terms of their relationship with the response variable. This can lead to an inconsistent result of variable importance when predictors are highly correlated (Strobl et al. 2008). In order to overcome this limitation, we repeated the analysis with conditional forest models using the “cforest” function (Strobl et al. 2009) in the package party in the R environment (Hothorn et al. 2006). Conditional forest models, similar to random forests, use an ensemble method that combines collection of trees, but use conditional inference trees instead of a regression tree and run a permutation test that is conditioned on the correlated predictors and ensure unbiased variable selection and variable importance. Since the conditional forest model results had very similar variable importance to random forest models, we continued with random forest model analyses.

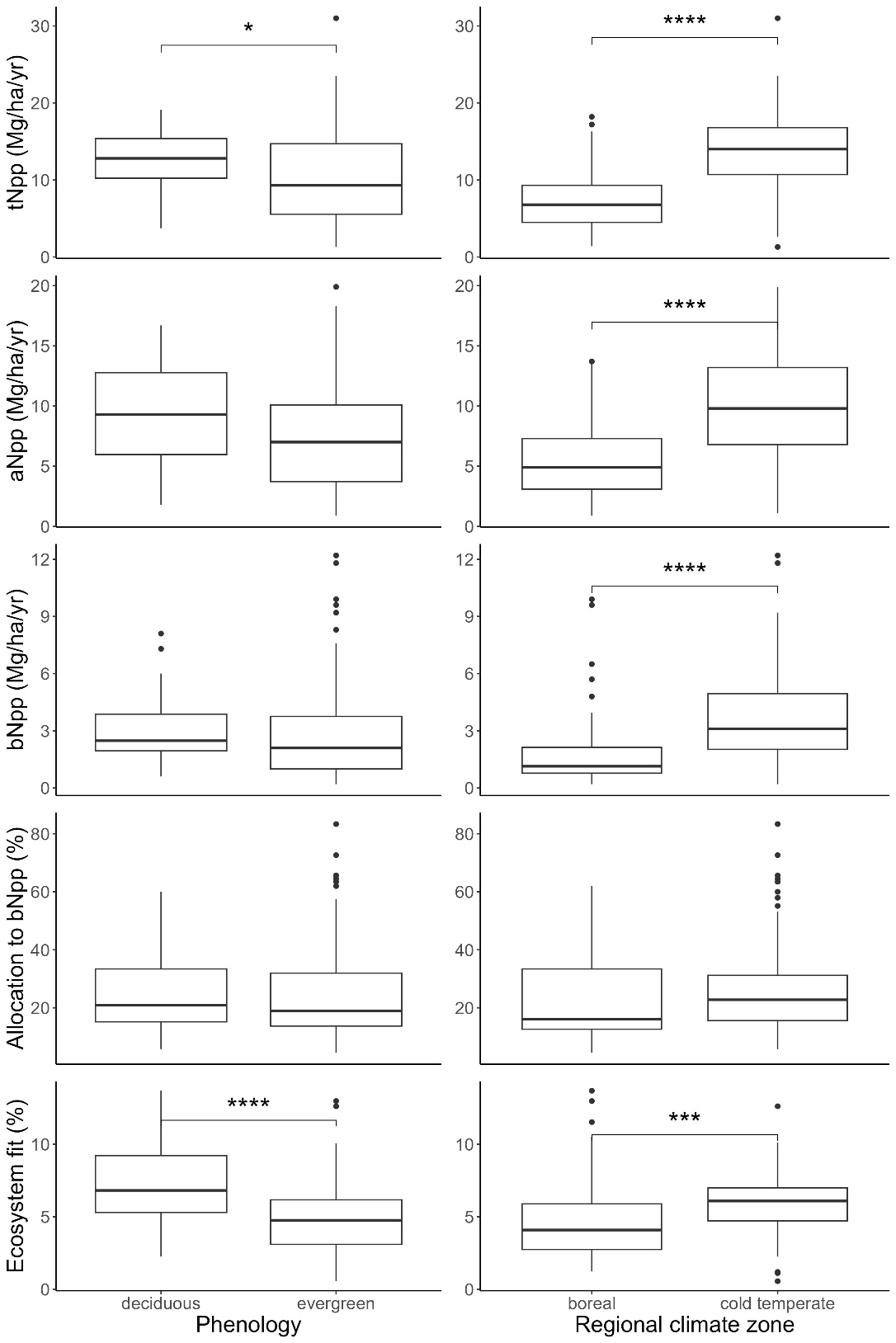

a.

b.

c.

d.

e.

**Fig. S1.** Comparison of (a) Total Net Primary Productivity (tNpp), (b) Aboveground Net Primary Productivity (aNpp), (c) Belowground Net Primary Productivity (bNpp), (d) fraction allocated belowground (% allocation to bNpp or bNpp/tNpp), and (e) ecosystem fit (a measure of theoretical productivity potential) divided by phenology (deciduous or evergreen; left column) and climate type (boreal, cold temperate, and warm temperate; right column). Statistics reported at top are Wilcoxon rank sum test and p value. Significant pairwise differences are reported with brackets for each pair (* p ≤ 0.05, ** p ≤ 0.01, *** p ≤ 0.001, **** p ≤ 0.0001)

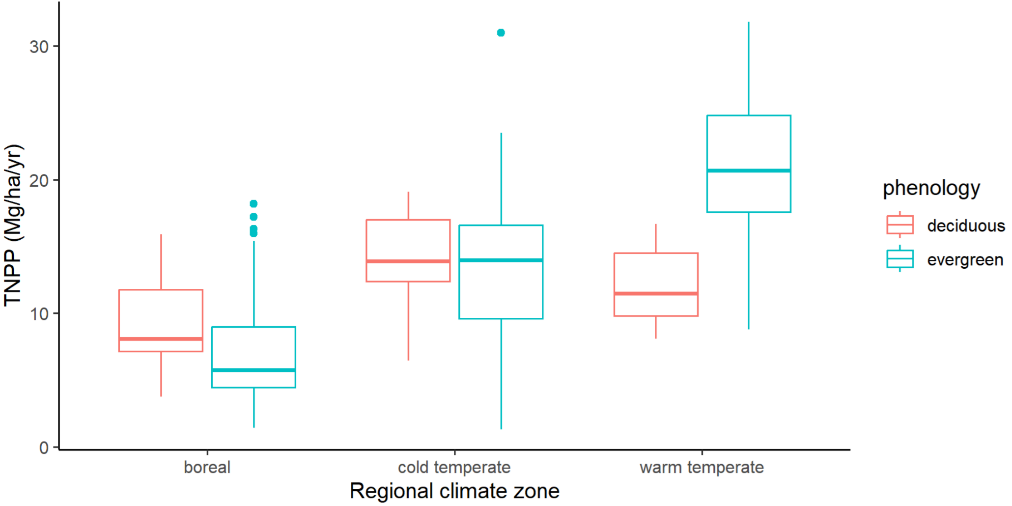

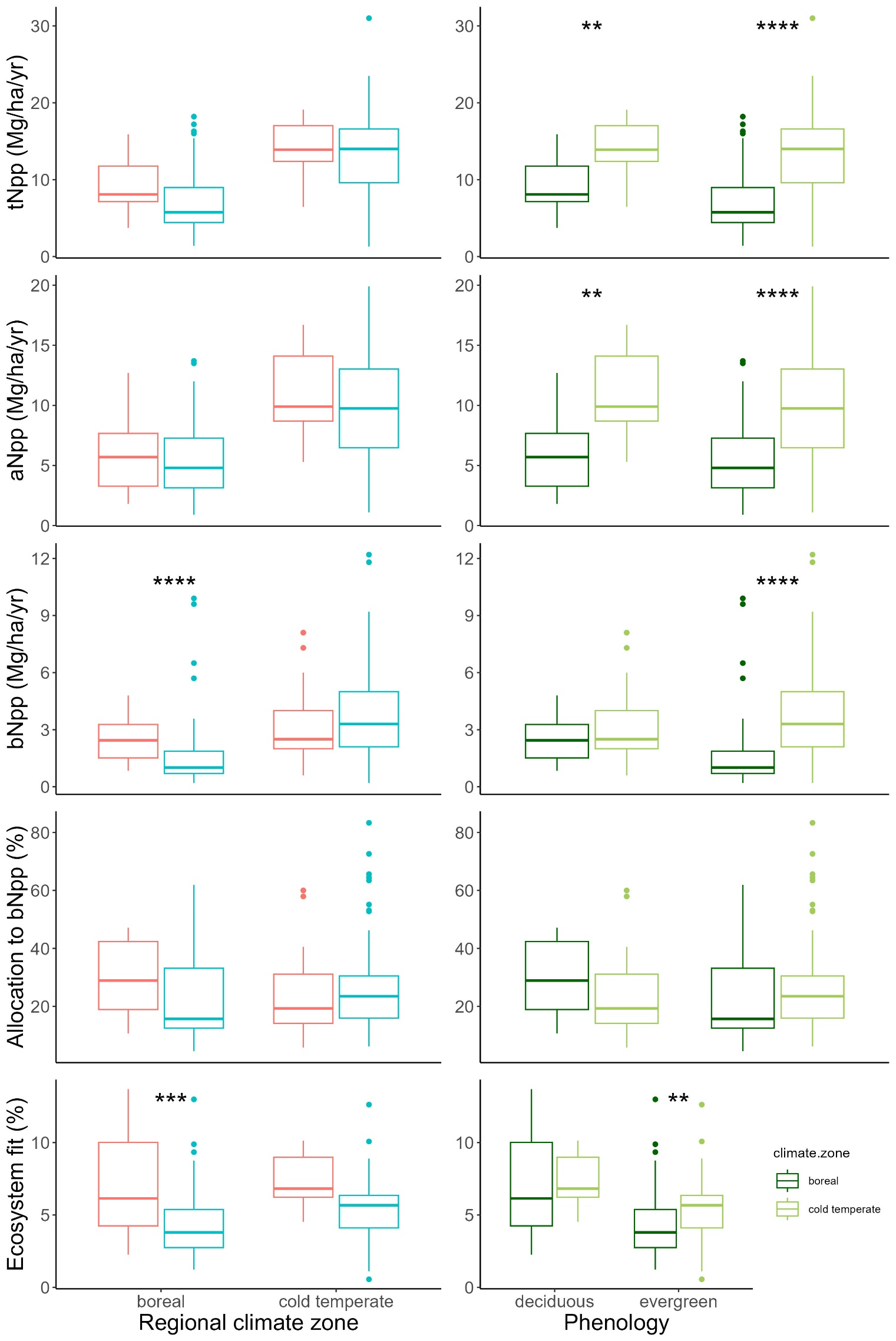

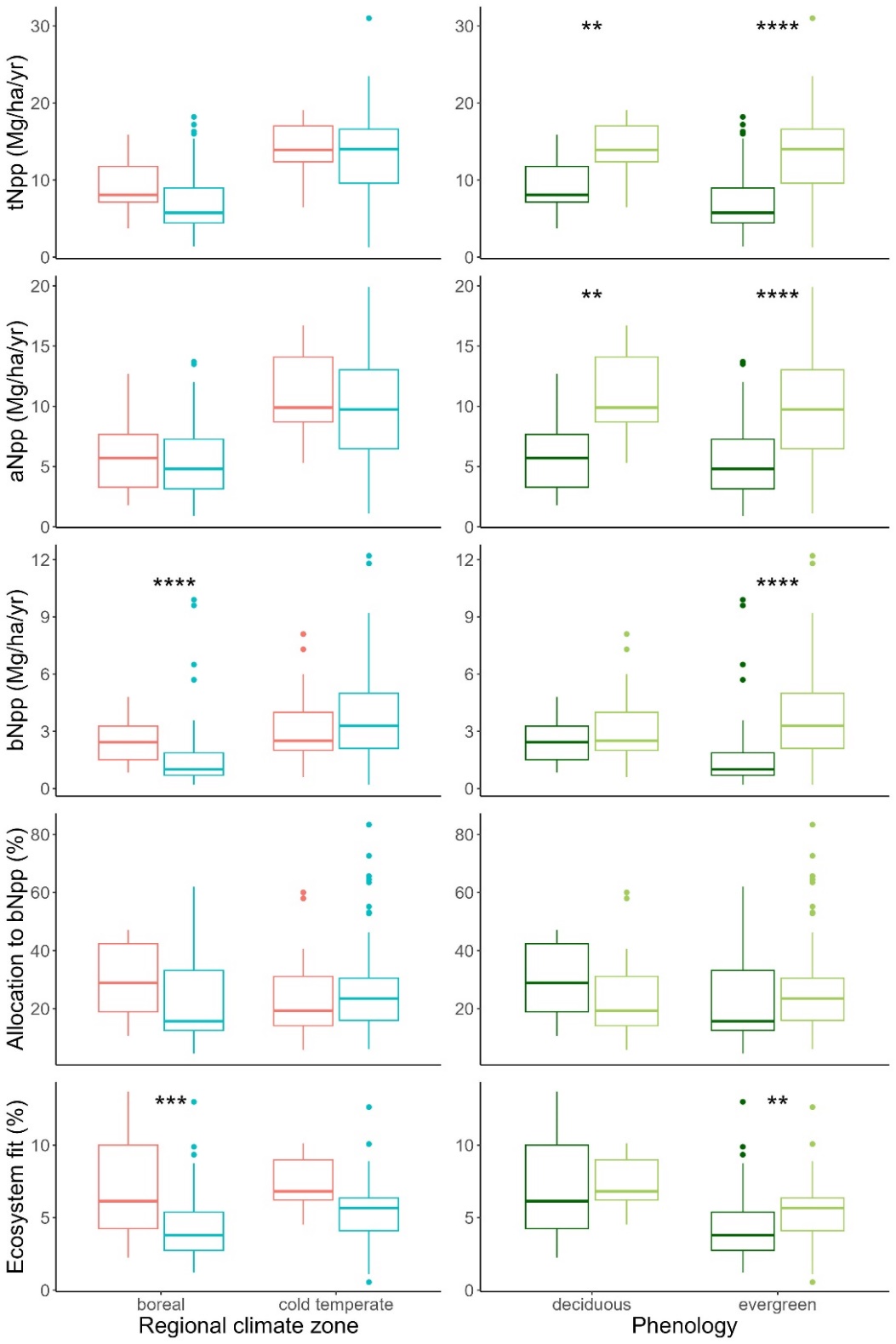

**Figure S2.** Comparison of (a) Total Net Primary Productivity (tNpp), (b) Aboveground Net Primary Productivity (aNpp), (c) Belowground Net Primary Productivity (bNpp), (d) fraction allocated belowground (% bNpp or bNpp/tNpp) and (e) ecosystem fit (a measure of theoretical productivity potential) divided by phenology (deciduous or evergreen) within climate type (boreal, cold temperate, and warm temperate; left column), and by climate type within phenology (right column). Statistics reported at top are Wilcoxon rank sum test and p value. Significant pairwise differences are reported with brackets for each pair (* p ≤ 0.05, ** p ≤ 0.01, *** p ≤ 0.001, **** p ≤ 0.0001).

B.

A.

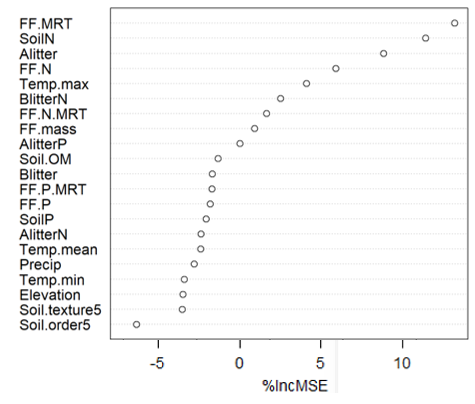

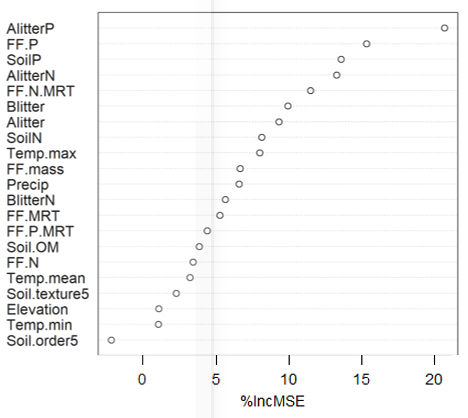

**Figure S3.** Variable importance plots generated from random forest regression modeling the effect of climatic and edaphic variables on percent allocation belowground (% bNpp) for boreal forests for A) low values of percent allocation belowground (< 20%) and B) high values of percent allocation belowground (> 20%).

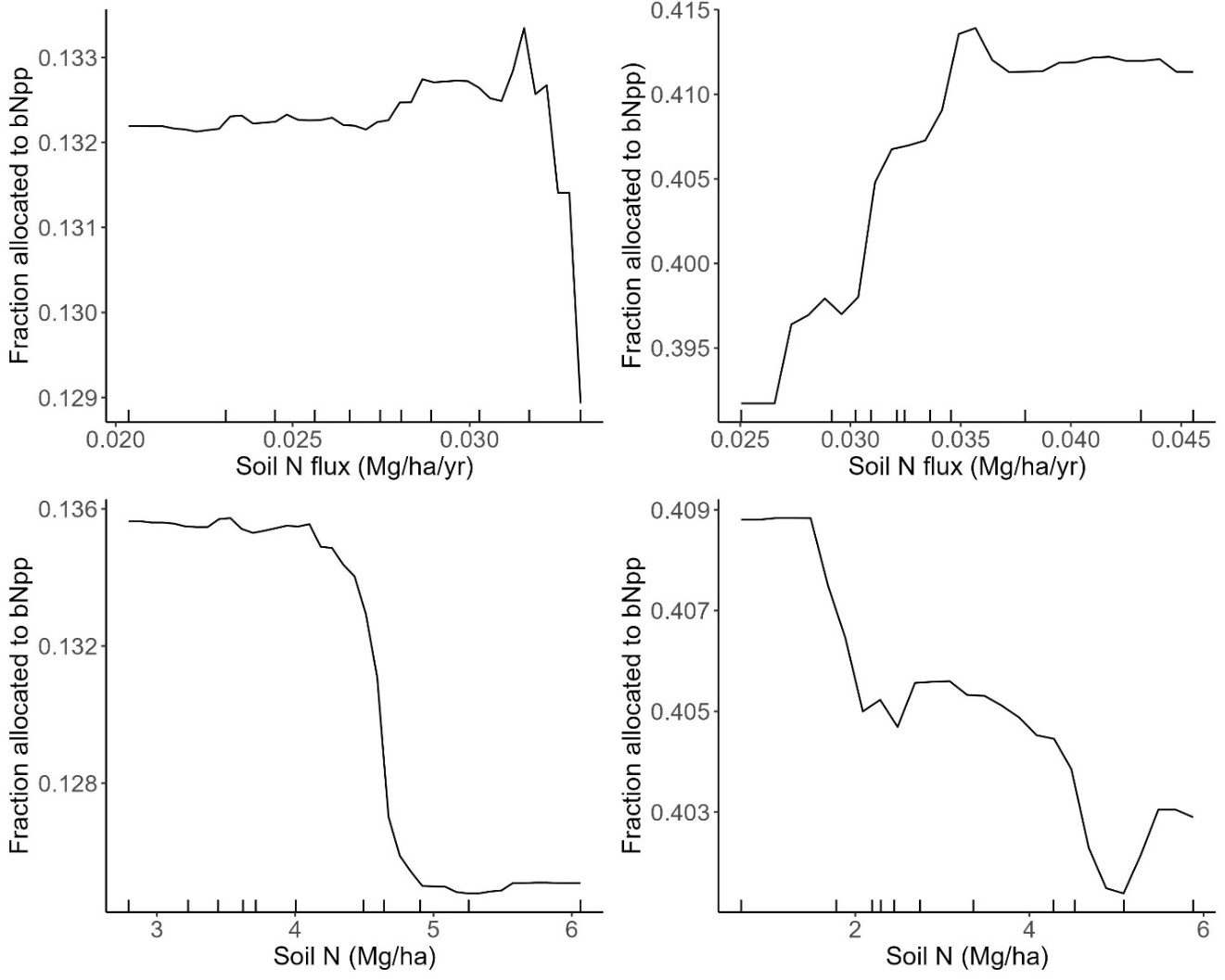

A.

B.

C.

D.

**Figure S4.** Partial dependence plots illustrating the effects of belowground litter N flux and the soil N pool on (A-B) low allocation (% bNpp < 20%) boreal forest sites and (C-D) high allocation (% bNpp > 20%) boreal forest sites.

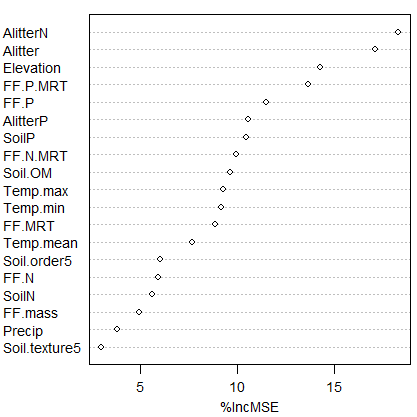

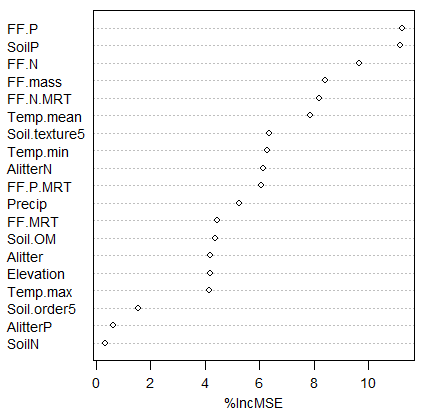

A.

B.

**Figure S5.** Variable importance plots generated from random forest regression modeling the effect of climatic and edaphic variables on belowground productivity (bNpp) for boreal forests for A) low values of percent allocation belowground (<2 Mg/ha/yr) and B) high values of percent allocation belowground (>2 Mg/ha/yr).

**Table S1.** Description of response variables and predictor variables used for this study. (Total Net Primary Productivity (tNpp), Above Net Primary Productivity (aNpp) and Belowground Net Primary Productivity (bNpp)).

| Response Variables | | Predictor Variables | | | | |
| --- | --- | --- | --- | --- | --- | --- |
| Variable Name | Description | Climate and elevation | Soil | | | |
|  |  |  | Soil type | Soil pools (Mg/ha) | Soil fluxes (Mg/ha/yr) | Soil fluxes (yrs) |
| aNpp (Mg/ha/yr) | Aboveground net primary productivity | Mean air temperature (C) | Soil taxonomic order | Soil N | Aboveground litter flux | Forest floor mean residence time |
| bNpp (Mg/ha/yr) | Belowground net primary productivity | Minimum air temperature (C) | Soil texture | Soil P | Belowground litter flux | Forest floor N mean residence time |
| tNpp (Mg/ha/yr)  [ tNpp =aNpp+bNpp] | Total net primary productivity | Maximum air temperature (C) |  | Soil organic matter | Aboveground litter N flux | Forest floor P mean residence time |
| % allocation to bNpp [bNpp/tNpp*100] | % tNpp allocated belowground | Total annual precipitation (mm) |  | Forest floor mass | Aboveground litter P flux |  |
| TMNPP (Mg/ha/yr)  [solar radiation * photosynthetic efficiency * interception efficiency * inverse of biomass * absorption efficiency] | Theoretical maximum net primary productivity | Elevation (m) |  | Forest floor N | Soil N flux |  |
| Ecosystem fit | tNpp/tmNpp |  |  | Forest floor P |  |  |

Table S2. Model summary for predicting tNpp, bNpp, and % allocation to bNpp for boreal and cold temperate forests. Pseudo R^2^ is the percent variance explained by out-of-bag (OOB) predictions. MSE OOB error is the mean squared error of out-of-bag errors. Model root mean square error (RMSE) is the tuned model error rate in units of the response. Mean absolute error (MAE) is the error rate for the model robustness against outliers.

| Model | Pseudo R^2^ | MSE OOB error | Model RMSE | MAE |
| --- | --- | --- | --- | --- |
| Boreal forests (n = 69):  tNpp  bNpp  % Allocation to bNpp | 73.2  74.51  68.9 | 4.2  0.9  0.007 | 0.89  0.44  0.03 | 0.71  0.27  0.02 |
| Cold temperate forests (n = 99):  tNpp  bNpp  % Allocation to bNpp | 64.17  45.13  44.06 | 8.44  2.74  0.01 | 1.25  0.67  0.05 | 0.85  0.49  0.03 |

**Table S3.** The most important predictor variables and their effect on response variables - TNPP, ANPP, BNPP, % BNPP, and ecosystem fit - by forest type (boreal, cold temperate, and warm temperate). Effects include trend (direction of effect), trend type (monotonic, non-monotonic, or hard saturation), threshold and saturating values for predictors, and the range of effect that each predictor had on the response variable.

| **TNPP** | | | | | |
| --- | --- | --- | --- | --- | --- |
| Climate | Variable | Trend | Trend type | Threshold & saturating values (predictor) | Range of effect on TNPP (Mg/ha/yr) |
| Boreal (n=69) | AlitterN (Mg/ha/yr) | + | m | 0.02-0.035 | 6.5-9.5 |
|  | FF.P.MRT (yrs) | - | m | 15-80 | 6.5-8.3 |
|  | Alitter (Mg /ha/yr) | + | m | 1.5-1.6,  2.5-5 | 6.5-8 |
| Cold temperate (n=99) | Soil N flux (Mg/ha/yr) | + | hs | 0.02-0.037 | 13-15.5 |
|  | AlitterP (Mg/ha/yr) | + | m | 0.002-0.004 | 12.5-15.5 |
|  | FF.P.MRT (yrs) | - | m | 0-60 | 12-14.5 |
|  | FF.MRT (yrs) | - | (hs) | 0-50 | 12.5-14 |

| **ANPP** | | | | | |
| --- | --- | --- | --- | --- | --- |
| Climate | Variable | Trend | Trend type | Threshold to saturating value (predictor) | Range of effect on ANPP (Mg/ha/yr) |
| Boreal (n=69) | AlitterN (Mg/ha/yr) | + | m | 0.02-0.04 | 5-7.5 |
|  | Soil P (Mg/ha) | + | m | 0.4-1.1 | 5-6.5 |
|  | FF.P.MRT (yr) | - | m | 10-60 | 5-6 |
|  | FF.N.MRT (yr) | - | M,hs | 10-30, 70-100 | 5-6.2 |
| Cold temperate (n=99) | AlitterP (Mg/ha/yr) | + | m | 0.002-0.004 | 8.5-11.5 |
|  | FF.MRT (yr) | - | (m) | 30-50 | 8.5-10.5 |
|  | FF.N.MRT (yr) | - | m | 30-70 | 9.4-10.4 |

| **BNPP** | | | | | |
| --- | --- | --- | --- | --- | --- |
| Climate | Variable | Trend | Trend type | Threshold to saturating value (predictor) | Range of effect on BNPP (Mg/ha/yr) |
| Boreal (n=69) | FF.P (Mg/ha) | - | hs | 0.05-0.1 | 1.5-3.5 |
|  | Soil N flux (Mg/ha/yr) | + | m | 0.03-0.04 | 1.5-3.5 |
|  | FF.N (Mg/ha) | - | hs | 0.4-0.6 | 1.8-2.8 |
| Cold temperate (n=99) | Soil N flux (Mg/ha/yr) | + | m | 0.02-0.09 | 3.5-5.5 |
|  | FF.P (Mg/ha) | - | m | 0-0.07 | 2.5-5.5 |

| **Percent BNPP** | | | | | |
| --- | --- | --- | --- | --- | --- |
| Climate | Variable | Trend | Trend type | Predictor: threshold and/or range of effect | Response: Range of effect |
| Boreal (n=69) | Soil N flux (Mg/ha/yr) | + | hs | 0.031 | 20-40 |
|  | Soil N (Mg/ha) | - | hs | 2.61 | 22-32 |
| Cold temperate (n=99) | Soil N flux (Mg/ha/yr) | + | hs,m | 4, 3-10 | 25-30,  30-33 |
|  | FF.N.MRT (yr) | -,+ | nm | 0-25 (-)  25-75(+) | 26-28.5 |
|  | AlitterP (Mg/ha/yr) | - | hs,m | 0.002,  0.002-0.004 | 30.5-26.5  25-26.5 |
|  | FF.P (Mg/ha) | - | m | 0-0.1 | 26-29 |
|  | FF.MRT (yr) | -,+ | nm | 0-10 (-),  10-60 (+) | 26.5-28.5 |
|  | Soil N flux (Mg/ha/yr) | -,+ | nm | 0.01-0.02 (-),  0.02-0.1 (+) | 26-29 |

| **Ecosystem Fit** | | | | | |
| --- | --- | --- | --- | --- | --- |
| Climate | Variable | Trend | Trend type | Threshold to saturating value (predictor) | Range of effect on ecosystem fit |
| Boreal (n=69) | AlitterN (Mg/ha/yr) | + | m | 0.02-0.035 | 0.045-0.058 |
|  | FF.P.MRT (yr) | - | m | No threshold, 10-80 | 0.044-0.054 |
|  | Alitter (Mg/ha/yr) | + | m | 1.5-5 | 0.045-0.051 |
| Cold temperate (n=99) | FF.P.MRT (yr) | - | m | 20-60 | 0.048-0.064 |
|  | Alitter P (Mg/ha/yr) | + | m | 0.002-0.004 | 0.05-0.075 |

**Table S4.** Ranked clusters of bNpp (belowground net primary productivity) and % allocation to bNpp (low, medium, and high) by ecophysiological scale. Notation shows mean ± SEM, [range], with number of samples (n) within each cluster. Clusters were selected using the partitioning around medoids (PAM) method. All bNpp and % allocation to bNpp mean values in the low, medium, and high ranks are significantly different within each row.

| **BNPP** | | | |
| --- | --- | --- | --- |
|  | Low | Medium | High |
| **Boreal and temperate (n=168)** | 1.1 ± 0.06 [0.2-2.1] *n*=77 | 3.2 ± 0.09 [2.2-4.8] *n*=61 | 6.8 ± 3.6 [5-12.2] *n*=30 |
| **Boreal (n=69)** | 0.93 ± 0.6 [0.2-1.9]  *n* =47 | 2.9 ± 0.2 [2-5.1]  *n*=18 | 7.9 ± 1 [5.7-9.9]  *n*=4 |
| **Cold temperate (n=99)** | 2.1 ± 0.1 [0.2-3.4] *n*=57 | 4.9 ± 0.2 [3.6-6.5] *n*=35 | 9.2 ± 0.8 [7.3-12.2] *n*=7 |
| **Evergreen (n=139)** | 1.4 ± 0.08 [0.2-2.8] *n*=85 | 4.5 ± 0.18 [3-6.5] *n*=45 | 9.8 ± 0.49 [7.6-12.2] *n*=9 |
| **Deciduous (n=42)** | 1.5 ± 0.1 [0.6-2.1] *n*=14 | 3 ± 0.1 [2.3-4]  *n*=18 | 6.2 ± 0.4 [4.8-8.1] *n*=10 |

| **% allocation to BNPP** | | | |
| --- | --- | --- | --- |
|  | Low | Medium | High |
| **Boreal and temperate (n=168)** | 15 ± 0.4 [4.5-23] *n*=92 | 31 ± 0.8 [23-43]  *n*=53 | 55 ± 2 [43-84]  *n*=23 |
| **Boreal (n=69)** | 13 ± 0.5 [4.5-20] *n*=42 | 31 ± 0.8 [26-35]  *n*=11 | 48 ± 1.5 [40-62]  *n*=16 |
| **Cold temperate (n=99)** | 17 ± 0.6 [5.8-24.7] *n*=59 | 33 ± 0.9 [25-46]  *n*=30 | 63 ± 3 [52-83]  *n*=10 |
| **Evergreen (n=139)** | 14 ± 0.4 [3.2-21] *n*=71 | 30 ± 0.8 [22-43]  *n*=49 | 57 ± 2.4 [44-83]  *n*=19 |
| **Deciduous (n=42)** | 16 ± 0.8 [6-21]  *n*=22 | 29 ± 1.3 [23-37]  *n*=10 | 48 ± 2.1 [39-60]  *n*=10 |

**Table S5.** Model comparison between linear regression, conditional forest (Cforest), and random forest models predicting Total Net Primary Productivity (tNpp), Above Net Primary Productivity (aNpp) and Belowground Net Primary Productivity (bNpp) for climate and edaphic variables using random mean squared error (RMSE) and mean absolute error (MAE).

|  | **tNpp** | |
| --- | --- | --- |
| **Model type** | **RMSE** | **MAE** |
| Linear regression | 3.3 | 2.6 |
| Cforest | 1.77 | 2.46 |
| Random forest | 0.04 | 1.08 |

|  | **aNpp** | |
| --- | --- | --- |
| **Model type** | **RMSE** | **MAE** |
| Linear regression | 2.96 | 2.24 |
| Cforest | 6.75 | 1.7 |
| Random forest | 1.04 | 0.74 |

|  | **bNpp** | |
| --- | --- | --- |
| **Model type** | **RMSE** | **MAE** |
| Linear regression | 1.6 | 1.14 |
| Cforest | 1.43 | 0.88 |
| Random forest | 0.58 | 0.39 |

**Table S6.** Database ID, site location, dominant forest type and citations for boreal and cold temperate forests.

| ID | Location | Forest Type | Citations |
| --- | --- | --- | --- |
| BOREAL FORESTS | | | |
| 1 | Alaska | *Picea mariana*, feather moss, no permafrost | Cole DW, M Rapp. 1981. Elemental cycling in forests. In: *Dynamic Properties of Forest Ecosystems* (ed Reichle DE), pp. 341-409. International Biological Programme 23, Cambridge Univ. Press, London; DeAngelis DL, RH Gardner, HH Shugart. 1981. Productivity of forest ecosystems studies during the IBP: the woodlands data set. In: Dynamic Properties of Forest Ecosystems (DE Reichle,ed) pp. 567-672. Cambridge University Press, Cambridge, UK; Soil Survey of Greater Nenana Area, Alaska. MRCS. <https://www.nrcs.usda.gov/Internet/FSE_MANUSCRIPTS/alaska/AK655/0/GreaterNenana.pdf>; Viereck LA, CT Dyrness, K Van Cleve, M J Foote. 1983. Vegetation, soils, and forest productivity in selected forest types in interior Alaska. Can. J. Forest Res. 13(5), 703-720. |
| 2 | Alaska | *Picea mariana*, muskeg, no permafrost | Cole DW, M Rapp. 1981. Elemental cycling in forests. In: *Dynamic Properties of Forest Ecosystems* (ed Reichle DE), pp. 341-409. International Biological Programme 23, Cambridge Univ. Press, London; DeAngelis DL, RH Gardner, HH Shugart. 1981. Productivity of forest ecosystems studies during the IBP: the woodlands data set. In: Dynamic Properties of Forest Ecosystems (DE Reichle,ed) pp. 567-672. Cambridge University Press, Cambridge, UK; Soil Survey of Greater Nenana Area, Alaska. MRCS. <https://www.nrcs.usda.gov/Internet/FSE_MANUSCRIPTS/alaska/AK655/0/GreaterNenana.pdf>; Viereck LA, CT Dyrness, K Van Cleve, M J Foote. 1983. Vegetation, soils, and forest productivity in selected forest types in interior Alaska. Can. J. Forest Res. 13(5), 703-720. |
| 3 | Alaska | *Picea mariana*, muskeg, permafrost 55 cm | Cole DW, M Rapp. 1981. Elemental cycling in forests. In: *Dynamic Properties of Forest Ecosystems* (ed Reichle DE), pp. 341-409. International Biological Programme 23, Cambridge Univ. Press, London; DeAngelis DL, RH Gardner, HH Shugart. 1981. Productivity of forest ecosystems studies during the IBP: the woodlands data set. In: Dynamic Properties of Forest Ecosystems (DE Reichle,ed) pp. 567-672. Cambridge University Press, Cambridge, UK; Soil Survey of Greater Nenana Area, Alaska. MRCS. <https://www.nrcs.usda.gov/Internet/FSE_MANUSCRIPTS/alaska/AK655/0/GreaterNenana.pdf>; Viereck LA, CT Dyrness, K Van Cleve, M J Foote. 1983. Vegetation, soils, and forest productivity in selected forest types in interior Alaska. Can. J. Forest Res. 13(5), 703-720. |
| 4 | Alaska, US [Bonanza Creek Experimental Forest LTER] | *Betula papyrifera* | Gower ST, O Krankina, RJ Olson, M Apps, S Linder, C Wang. 2001. Net primary production and carbon allocation patterns of boreal forest ecosystems. Ecol. Appl 11(5), 1395-1411; Soil Survey of Greater Nenana Area, Alaska. MRCS. <https://www.nrcs.usda.gov/Internet/FSE_MANUSCRIPTS/alaska/AK655/0/GreaterNenana.pdf>; Van Cleve K, LA Viereck, CT Dyrness. 1996. State factor control of soils and forest succession along the Tanana River in interior Alaska, U.S.A. Arctic and Alpine Research 28:388-400. |
| 5 | Alaska | *Betula papyrifera* | Cole DW, M Rapp. 1981. Elemental cycling in forests. In: *Dynamic Properties of Forest Ecosystems* (ed Reichle DE), pp. 341-409. International Biological Programme 23, Cambridge Univ. Press, London; Soil Survey of Greater Nenana Area, Alaska. MRCS. <https://www.nrcs.usda.gov/Internet/FSE_MANUSCRIPTS/alaska/AK655/0/GreaterNenana.pdf>; Van Cleve K, LA Viereck, and CT Dyrness. 1996. State factor control of soils and forest succession along the Tanana River in interior Alaska, U.S.A. Arctic and Alpine Research 28:388-400; Viereck LA, CT Dyrness, K Van Cleve, MJ Foote. 1983. Vegetation, soils, and forest productivity in selected forest types in interior Alaska. Can. J. Forest Res. 13(5), 703-720. |
| 6 | Alaska | *Picea glauca* | Gower ST, O Krankina, RJ Olson, M Apps, S Linder, C Wang. 2001. Net primary production and carbon allocation patterns of boreal forest ecosystems. Ecol. Appl 11(5), 1395-1411; Soil Survey of Greater Nenana Area, Alaska. MRCS. https://www.nrcs.usda.gov/Internet/FSE_MANUSCRIPTS/alaska/AK655/0/GreaterNenana.pdf |
| 7 | Alaska | *Picea glauca* | Gower ST, O Krankina, RJ Olson, M Apps, S Linder, C Wang. 2001. Net primary production and carbon allocation patterns of boreal forest ecosystems. Ecol. Appl 11(5), 1395-1411; Soil Survey of Greater Nenana Area, Alaska. MRCS. <https://www.nrcs.usda.gov/Internet/FSE_MANUSCRIPTS/alaska/AK655/0/GreaterNenana.pdf>; Viereck LA, K Van Cleve, PC Adams, RE Schlentner. 1993. Climate of the Tanana River floodplain near Fairbanks, Alaska. CJFR 23, 899-913 |
| 8 | Alaska | *Picea mariana* - black spruce | Gower ST, O Krankina, RJ Olson, M Apps, S Linder, C Wang. 2001. Net primary production and carbon allocation patterns of boreal forest ecosystems. Ecol. Appl 11(5), 1395-1411; Soil Survey of Greater Nenana Area, Alaska. MRCS. <https://www.nrcs.usda.gov/Internet/FSE_MANUSCRIPTS/alaska/AK655/0/GreaterNenana.pdf>; Viereck LA, K Van Cleve, PC Adams, RE Schlentner. 1993. Climate of the Tanana River floodplain near Fairbanks, Alaska. CJFR 23, 899-913 |
| 9 | Alaska | *Populus balsamifera* | Gower ST, O Krankina, RJ Olson, M Apps, S Linder, C Wang. 2001. Net primary production and carbon allocation patterns of boreal forest ecosystems. Ecol. Appl 11(5), 1395-1411; Soil Survey of Greater Nenana Area, Alaska. MRCS. https://www.nrcs.usda.gov/Internet/FSE_MANUSCRIPTS/alaska/AK655/0/GreaterNenana.pdf |
| 10 | Alaska | *Populus/Alnus* | Gower ST, O Krankina, RJ Olson, M Apps, S Linder, C Wang. 2001. Net primary production and carbon allocation patterns of boreal forest ecosystems. Ecol. Appl 11(5), 1395-1411; Soil Survey of Greater Nenana Area, Alaska. MRCS. https://www.nrcs.usda.gov/Internet/FSE_MANUSCRIPTS/alaska/AK655/0/GreaterNenana.pdf |
| 11 | Canada, Manitoba [BOREAS NSA] | *Picea mariana* | Gower ST, O Krankina, RJ Olson, M Apps, S Linder, C Wang. 2001. Net primary production and carbon allocation patterns of boreal forest ecosystems. Ecol. Appl 11(5), 1395-1411; Gower ST, JG Vogel, JM Norman, CJ Kucharik, SJ Steele, TK Stow. 1997. Carbon distribution and aboveground net primary production in aspen, jack pine, and black spruce stands in Saskatchewan and Manitoba, Canada. J Geophysical Res 102, 29029-29041 |
| 12 | Canada, Manitoba [BOREAS NSA] | *Pinus banksiana -* Jack pine | Gower ST, O Krankina, RJ Olson, M Apps, S Linder, C Wang. 2001. Net primary production and carbon allocation patterns of boreal forest ecosystems. Ecol. Appl 11(5), 1395-1411; Gower ST, JG Vogel, JM Norman, CJ Kucharik, SJ Steele, TK Stow. 1997. Carbon distribution and aboveground net primary production in aspen, jack pine, and black spruce stands in Saskatchewan and Manitoba, Canada. Journal of Geophysical Res 102: 29,029-29,041 |
| 13 | Canada, Manitoba [BOREAS NSA] | *Populus tremuloides* - trembling aspen | Gower ST, JG Vogel, JM Norman, CJ Kucharik, SJ Steele, TK Stow. 1997. Carbon distribution and aboveground net primary production in aspen, jack pine, and black spruce stands in Saskatchewan and Manitoba, Canada. Journal of Geophysical Res 102: 29,029-29,041; Gower ST, O Krankina, RJ Olson, M Apps, S Linder, C Wang. 2001. Net primary production and carbon allocation patterns of boreal forest ecosystems. Ecol. Appl 11(5), 1395-1411 |
| 14 | Canada, Ontario | *Picea rubens*, dry | DeAngelis DL, RH Gardner, HH Shugart. 1981. Productivity of forest ecosystems studies during the IBP: the woodlands data set. In: Dynamic Properties of Forest Ecosystems (DE Reichle,ed) pp. 567-672. Cambridge University Press, Cambridge, UK; Van Cleve K, LA Viereck, CT Dyrness. 1996. State factor control of soils and forest succession along the Tanana River in interior Alaska, U.S.A. Arctic and Alpine Research 28:388-400. |
| 15 | Canada, Ontario | *Picea rubens*, fresh | DeAngelis DL, RH Gardner, HH Shugart. 1981. Productivity of forest ecosystems studies during the IBP: the woodlands data set. In: Dynamic Properties of Forest Ecosystems (DE Reichle,ed) pp. 567-672. Cambridge University Press, Cambridge, UK. |
| 16 | Canada, Ontario | *Picea rubens*, moist | DeAngelis DL, RH Gardner, HH Shugart. 1981. Productivity of forest ecosystems studies during the IBP: the woodlands data set. In: Dynamic Properties of Forest Ecosystems (DE Reichle,ed) pp. 567-672. Cambridge University Press, Cambridge, UK. |
| 17 | Canada, Ontario | *Picea rubens*, wet | DeAngelis DL, RH Gardner, HH Shugart. 1981. Productivity of forest ecosystems studies during the IBP: the woodlands data set. In: Dynamic Properties of Forest Ecosystems (DE Reichle,ed) pp. 567-672. Cambridge University Press, Cambridge, UK. |
| 18 | Canada, Saskatchewan [BOREAS SSA] | *Picea mariana* - black spruce | Gower ST, JG Vogel, JM Norman, CJ Kucharik, SJ Steele, TK Stow. 1997. Carbon distribution and aboveground net primary production in aspen, jack pine, and black spruce stands in Saskatchewan and Manitoba, Canada. Journal of Geophysical Res 102, 29029-29041; Gower ST, O Krankina, RJ Olson, M Apps, S Linder, C Wang. 2001. Net primary production and carbon allocation patterns of boreal forest ecosystems. Ecol. Appl 11(5), 1395-1411; Van Cleve K, LA Viereck, CT Dyrness. 1996. State factor control of soils and forest succession along the Tanana River in interior Alaska, U.S.A. Arctic and Alpine Research 28, 388-400. |
| 19 | Canada, Saskatchewan [BOREAS SSA] | *Picea mariana* - closed canopy feathermoss ground cover (BSFM) | Gower ST, JG Vogel, JM Norman, CJ Kucharik, SJ Steele, TK Stow. 1997. Carbon distribution and aboveground net primary production in aspen, jack pine, and black spruce stands in Saskatchewan and Manitoba, Canada. Journal of Geophysical Res 102, 29029-29041; Gower ST, O Krankina, RJ Olson, M Apps, S Linder, C Wang. 2001. Net primary production and carbon allocation patterns of boreal forest ecosystems. Ecol. Appl 11(5), 1395-1411; O'Connell KEB, Gower ST, Norman JM. 2003. Comparison of Net Primary Production and light-use dynamics of two boreal black spruce forest communities. Ecosystems 6, 236-247. |
| 20 | Canada, Saskatchewan [BOREAS SSA] | *Picea mariana* - open canopy with Sphagnum ground cover (BSSP) | Gower ST, JG Vogel, JM Norman, CJ Kucharik, SJ Steele, TK Stow. 1997. Carbon distribution and aboveground net primary production in aspen, jack pine, and black spruce stands in Saskatchewan and Manitoba, Canada. Journal of Geophysical Res 102, 2929-29041; Gower ST, O Krankina, RJ Olson, M Apps, S Linder, C Wang. 2001. Net primary production and carbon allocation patterns of boreal forest ecosystems. Ecol. Appl 11(5), 1395-1411; O'Connell KEB, Gower ST, Norman JM. 2003. Comparison of Net Primary Production and light-use dynamics of two boreal black spruce forest communities. Ecosystems 6, 236-247. |
| 21 | Canada, Saskat-chewan [BOREAS SSA] | *Pinus banksiana* - Jack pine | Gower ST, JG Vogel, JM Norman, CJ Kucharik, SJ Steele, TK Stow. 1997. Carbon distribution and aboveground net primary production in aspen, jack pine, and black spruce stands in Saskatchewan and Manitoba, Canada. Journal of Geophysical Res 102, 29029-29041; Gower ST, O Krankina, RJ Olson, M Apps, S Linder, C Wang. 2001. Net primary production and carbon allocation patterns of boreal forest ecosystems. Ecol. Appl 11(5), 1395-1411 |
| 22 | Canada, Saskat-chewan [BOREAS SSA] | *Populus tremuloides* - trembling aspen | Gower ST, JG Vogel, JM Norman, CJ Kucharik, SJ Steele, TK Stow. 1997. Carbon distribution and aboveground net primary production in aspen, jack pine, and black spruce stands in Saskatchewan and Manitoba, Canada. Journal of Geophysical Res 102, 29029-29041; Gower ST, O Krankina, RJ Olson, M Apps, S Linder, C Wang. 2001. Net primary production and carbon allocation patterns of boreal forest ecosystems. Ecol. Appl 11(5), 1395-1411 |
| 23 | China, Tahe, Daxing'anling | *Larix gmelinii* | Gower ST, O Krankina, RJ Olson, M Apps, S Linder, C Wang. 2001. Net primary production and carbon allocation patterns of boreal forest ecosystems. Ecol. Appl 11(5), 1395-1411; Leuschner C, A Zach, G Moser, J Homeier, S Graefe, D Hertel, B Wittich, N Soethe, S Iost, M Röderstein, V Horna, K Wolf. 2013. The Carbon Balance of Tropical Mountain Forests Along an Altitudinal Transect. In: Bendix J. et al. (eds) Ecosystem Services, Biodiversity and Environmental Change in a Tropical Mountain Ecosystem of South Ecuador. Ecological Studies (Analysis and Synthesis), vol 221, pp. 117-139. Springer, Berlin, Heidelberg. <https://doi.org/10.1007/978-3-642-38137-9_10>; Wang C, ST Gower, Y Wang, H Zhao, P Yan, B. Bond-Lamberty. 2001. Influence of fire on carbon distribution and net primary production of boreal *Larix gmelinii* forests in northeastern China. Global Change Biol 7, 719– 730. |
| 24 | Finland | *Picea excelsa* | DeAngelis DL, RH Gardner, HH Shugart. 1981. Productivity of forest ecosystems studies during the IBP: the woodlands data set. In: Dynamic Properties of Forest Ecosystems (DE Reichle,ed) pp. 567-672. Cambridge University Press, Cambridge, UK; Kimmins JP, BC Hawkes. 1978. Distribution and chemistry of fine roots in a white spruce-subalpine fir stand in British Columbia: Implications for management. Can. J. For. Res. 8, 265-279. |
| 25 | Finland, Ilomantsi [RhNRmu], eastern Finland | *Betula pubescens* | Finér L. 1989. Biomass and nutrient cycle in fertilized and unfertilized pine, mixed birch and pine and spruce stands on a drained mire. Acta Forestalia Fennica 208, 1-63; Gower ST, O Krankina, RJ Olson, M Apps, S Linder, C Wang. 2001. Net primary production and carbon allocation patterns of boreal forest ecosystems. Ecol. Appl 11(5), 1395-1411 |
| 26 | Finland, Ilomantsi [Mkmu] | *Picea abies* | Finér L. 1989. Biomass and nutrient cycle in fertilized and unfertilized pine, mixed birch and pine and spruce stands on a drained mire. Acta Forestalia Fennica 208, 1-63; Gower ST, O Krankina, RJ Olson, M Apps, S Linder, C Wang. 2001. Net primary production and carbon allocation patterns of boreal forest ecosystems. Ecol. Appl 11(5), 1395-1411 |
| 27 | Finland, Ilomantsi [VNRmu] | *Pinus sylvestris* | Finér L. 1989. Biomass and nutrient cycle in fertilized and unfertilized pine, mixed birch and pine and spruce stands on a drained mire. Acta Forestalia Fennica 208, 1-63; Gower ST, O Krankina, RJ Olson, M Apps, S Linder, C Wang. 2001. Net primary production and carbon allocation patterns of boreal forest ecosystems. Ecol. Appl 11(5), 1395-1411 |
| 28 | Finland, Ilomantsi [RhNRmu] | *Pinus sylvestris* | Finér L. 1989. Biomass and nutrient cycle in fertilized and unfertilized pine, mixed birch and pine and spruce stands on a drained mire. Acta Forestalia Fennica 208, 1-63; Gower ST, O Krankina, RJ Olson, M Apps, S Linder, C Wang. 2001. Net primary production and carbon allocation patterns of boreal forest ecosystems. Ecol. Appl 11(5), 1395-1411; O'Connell KEB, Gower ST, Norman JM. 2003. Net ecosystem production of two contrasting boreal black spruce forest communities. Ecosystems 6, 248-260 |
| 29 | Finland, Kuusamo [Oulanka National Park] | *Picea abies* | Havas, P. 2013. NPP Boreal Forest: Kuusamo, Finland, 1967-1972, R1. Data set. Available on-line [http://daac.ornl.gov] from Oak Ridge National Laboratory Distributed Active Archive Center, Oak Ridge, Tennessee, USA doi:10.3334/ORNLDAAC/466; O'Connell KEB, ST Gower, JM Norman. 2003. Net ecosystem production of two contrasting boreal black spruce forest communities. Ecosystems 6, 248-260. |
| 30 | Iceland, Eastern [Hallormsstadur] | *Betula pubescen* | Sigurdardottir R. 1999. Effects of different forest types on total ecosystem carbon sequestration in Hallormsstadur Forest, Eastern Iceland. (Dissertation, Yale) |
| 31 | Iceland, Eastern [Hallormsstadur] | *Larix sibirica* | Sigurdardottir R. 1999. Effects of different forest types on total ecosystem carbon sequestration in Hallormsstadur Forest, Eastern Iceland. (Dissertation, Yale) |
| 32 | Iceland, Eastern [Hallormsstadur] | *Pinus contorta* | Sigurdardottir R. 1999. Effects of different forest types on total ecosystem carbon sequestration in Hallormsstadur Forest, Eastern Iceland. (Dissertation, Yale) |
| 33 | Russia (Siberia), Irkutsk | *Pinus sylvestris* | Krankina O.N. 2013. NPP Boreal Forest: Siberian Scots Pine Forests, Russia, 1968-1974 Revision 1. Data set. Available on-line [http://daac.ornl.gov] from Oak Ridge National Laboratory Distributed Active Archive Center, Oak Ridge, Tennessee, USA doi:10.3; Gower ST, O Krankina, RJ Olson, M Apps, S Linder, C Wang. 2001. Net primary production and carbon allocation patterns of boreal forest ecosystems. Ecol. Appl 11(5), 1395-1411 |
| 34 | Russia (Siberia), Irkutsk | *Pinus sylvestris* | Krankina O.N. 2013. NPP Boreal Forest: Siberian Scots Pine Forests, Russia, 1968-1974 Revision 1. Data set. Available on-line [http://daac.ornl.gov] from Oak Ridge National Laboratory Distributed Active Archive Center, Oak Ridge, Tennessee, USA doi:10.3; Gower ST, O Krankina, RJ Olson, M Apps, S Linder, C Wang. 2001. Net primary production and carbon allocation patterns of boreal forest ecosystems. Ecol. Appl 11(5), 1395-1411 |
| 35 | Russia (Siberia), Irkutsk | *Pinus sylvestris* | Krankina O.N. 2013. NPP Boreal Forest: Siberian Scots Pine Forests, Russia, 1968-1974 Revision 1. Data set. Available on-line [http://daac.ornl.gov] from Oak Ridge National Laboratory Distributed Active Archive Center, Oak Ridge, Tennessee, USA doi:10.3; Gower ST, O Krankina, RJ Olson, M Apps, S Linder, C Wang. 2001. Net primary production and carbon allocation patterns of boreal forest ecosystems. Ecol. Appl 11(5), 1395-1411 |
| 36 | Russia (Siberia), Tomsk (ssp1a) | *Pinus sylvestris* | Gower, S.T., O. Krankina, R.J. Olson, M. Apps, S. Linder, and C. Wang. 2012. NPP Boreal Forest: Consistent Worldwide Site Estimates, 1965-1995, R1. Data set. Available on-line [http://daac.ornl.gov] from the Oak Ridge National Laboratory Distributed Active Archive Center, Oak Ridge, Tennessee, U.S.A. |
| 37 | Russia (Siberia), Tomsk (ssp1b) | *Pinus sylvestris* | Gower, S.T., O. Krankina, R.J. Olson, M. Apps, S. Linder, C. Wang. 2012. NPP Boreal Forest: Consistent Worldwide Site Estimates, 1965-1995, R1. Data set. Available on-line [http://daac.ornl.gov] Oak Ridge National Laboratory Distributed Active Archive Center, Oak Ridge, Tennessee, U.S.A. |
| 38 | Russia (Siberia), Tomsk (ssp1c) | *Pinus sylvestris* | Gower, S.T., O. Krankina, R.J. Olson, M. Apps, S. Linder, and C. Wang. 2012. NPP Boreal Forest: Consistent Worldwide Site Estimates, 1965-1995, R1. Data set. Available on-line [http://daac.ornl.gov] Oak Ridge National Laboratory Distributed Active Archive Center, Oak Ridge, Tennessee, U.S.A. |
| 39 | Russia (Siberia), Tomsk (ssp1d) | *Pinus sylvestris* | Gower, S.T., O. Krankina, R.J. Olson, M. Apps, S. Linder, C. Wang. 2012. NPP Boreal Forest: Consistent Worldwide Site Estimates, 1965-1995, R1. Data set. Available on-line [http://daac.ornl.gov] Oak Ridge National Laboratory Distributed Active Archive Center, Oak Ridge, Tennessee, U.S.A. |
| 40 | Russia (Siberia), Tomsk (ssp1e) | *Pinus sylvestris* | Finér L. 1989. Biomass and nutrient cycle in fertilized and unfertilized pine, mixed birch and pine and spruce stands on a drained mire. Acta Forestalia Fennica 208, 1-63; Gower ST, O Krankina, RJ Olson, M Apps, S Linder, C Wang. 2001. Net primary production and carbon allocation patterns of boreal forest ecosystems. Ecol. Appl 11(5), 1395-1411 |
| 41 | Russia (Siberia), Tomsk (ssp1f) | *Pinus sylvestris* | Finér L. 1989. Biomass and nutrient cycle in fertilized and unfertilized pine, mixed birch and pine and spruce stands on a drained mire. Acta Forestalia Fennica 208, 1-63; Gower ST, O Krankina, RJ Olson, M Apps, S Linder, C Wang. 2001. Net primary production and carbon allocation patterns of boreal forest ecosystems. Ecol. Appl 11(5), 1395-1411 |
| 42 | Russia (Siberia), Tomsk (ssp1g) | *Pinus sylvestris* | Finér L. 1989. Biomass and nutrient cycle in fertilized and unfertilized pine, mixed birch and pine and spruce stands on a drained mire. Acta Forestalia Fennica 208, 1-63; Gower ST, O Krankina, RJ Olson, M Apps, S Linder, C Wang. 2001. Net primary production and carbon allocation patterns of boreal forest ecosystems. Ecol. Appl 11(5), 1395-1411 |
| 43 | Russia (Siberia), Tomsk (ssp1h) | *Pinus sylvestris* | Finér L. 1989. Biomass and nutrient cycle in fertilized and unfertilized pine, mixed birch and pine and spruce stands on a drained mire. Acta Forestalia Fennica 208, 1-63; Gower ST, O Krankina, RJ Olson, M Apps, S Linder, C Wang. 2001. Net primary production and carbon allocation patterns of boreal forest ecosystems. Ecol. Appl 11(5), 1395-1411 |
| 44 | Russia (Siberia), Tomsk (ssp1i) | *Pinus sylvestris* | Finér L. 1989. Biomass and nutrient cycle in fertilized and unfertilized pine, mixed birch and pine and spruce stands on a drained mire. Acta Forestalia Fennica 208, 1-63; Gower ST, O Krankina, RJ Olson, M Apps, S Linder, C Wang. 2001. Net primary production and carbon allocation patterns of boreal forest ecosystems. Ecol. Appl 11(5), 1395-1411 |
| 45 | Russia (Siberia), Tomsk (ssp1j) | *Pinus sylvestris* | Finér L. 1989. Biomass and nutrient cycle in fertilized and unfertilized pine, mixed birch and pine and spruce stands on a drained mire. Acta Forestalia Fennica 208, 1-63; Gower ST, O Krankina, RJ Olson, M Apps, S Linder, C Wang. 2001. Net primary production and carbon allocation patterns of boreal forest ecosystems. Ecol. Appl 11(5), 1395-1411 |
| 46 | Russia (Siberia), Tomsk (ssp1k) | *Pinus sylvestris* | Finér L. 1989. Biomass and nutrient cycle in fertilized and unfertilized pine, mixed birch and pine and spruce stands on a drained mire. Acta Forestalia Fennica 208, 1-63; Gower ST, O Krankina, RJ Olson, M Apps, S Linder, C Wang. 2001. Net primary production and carbon allocation patterns of boreal forest ecosystems. Ecol. Appl 11(5), 1395-1411 |
| 47 | Russia (Southern Karelia) (Site 1) | *Picea abies* - Southern Karelian spruce | DeAngelis DL, RH Gardner, HH Shugart. 1981. Productivity of forest ecosystems studies during the IBP: the woodlands data set. In: Dynamic Properties of Forest Ecosystems (DE Reichle, ed) pp. 567-672. Cambridge University Press, Cambridge, UK; Kazimirov NI, RM Morozova. 1973. Biological cycling of matter in spruce forests of Karelia. NAUKA, Leningrad Branch, Academy of Sciences. 168 pp. |
| 48 | Russia (Southern Karelia) (Site 10) | *Picea abies* - Southern Karelian spruce | Cole DW, M Rapp. 1981. Elemental cycling in forests. In: Dynamic Properties of Forest Ecosystems (DE Reichle, ed), pp. 341-409. International Biological Programme 23, Cambridge Univ. Press, London; DeAngelis DL, RH Gardner, HH Shugart. 1981. Productivity of forest ecosystems studies during the IBP: the woodlands data set. In: Dynamic Properties of Forest Ecosystems (DE Reichle, ed) pp. 567-672. Cambridge University Press, Cambridge, UK; Kazimirov NI, RM Morozova. 1973. Biological cycling of matter in spruce forests of Karelia. NAUKA, Leningrad Branch, Academy of Sciences. 168 pp. |
| 49 | Russia (Southern Karelia), (Site 11) | *Picea abies* - Southern Karelian spruce | DeAngelis DL, RH Gardner, HH Shugart. 1981. Productivity of forest ecosystems studies during the IBP: the woodlands data set. In: Dynamic Properties of Forest Ecosystems (DE Reichle, ed) pp. 567-672. Cambridge University Press, Cambridge, UK; Kazimirov NI, RM Morozova. 1973. Biological cycling of matter in spruce forests of Karelia. NAUKA, Leningrad Branch, Academy of Sciences. 168 pp. |
| 50 | Russia (Southern Karelia), (Site 12) | *Picea abies* - Southern Karelian spruce | DeAngelis DL, RH Gardner, HH Shugart. 1981. Productivity of forest ecosystems studies during the IBP: the woodlands data set. In: Dynamic Properties of Forest Ecosystems (DE Reichle, ed) pp. 567-672. Cambridge University Press, Cambridge, UK; Kazimirov NI, RM Morozova. 1973. Biological cycling of matter in spruce forests of Karelia. NAUKA, Leningrad Branch, Academy of Sciences. 168 pp. |
| 51 | Russia (Southern Karelia), (Site 13) | *Picea abies* - Southern Karelian spruce | DeAngelis DL, RH Gardner, HH Shugart. 1981. Productivity of forest ecosystems studies during the IBP: the woodlands data set. In: Dynamic Properties of Forest Ecosystems (DE Reichle, ed) pp. 567-672. Cambridge University Press, Cambridge, UK; Kazimirov NI, RM Morozova. 1973. Biological cycling of matter in spruce forests of Karelia. NAUKA, Leningrad Branch, Academy of Sciences. 168 pp. |
| 52 | Russia (Southern Karelia), (Site 14) | *Picea abies* - Southern Karelian spruce | DeAngelis DL, RH Gardner, HH Shugart. 1981. Productivity of forest ecosystems studies during the IBP: the woodlands data set. In: Dynamic Properties of Forest Ecosystems (DE Reichle, ed) pp. 567-672. Cambridge University Press, Cambridge, UK; Kazimirov NI, RM Morozova. 1973. Biological cycling of matter in spruce forests of Karelia. NAUKA, Leningrad Branch, Academy of Sciences. 168 pp. |
| 53 | Russia (Southern Karelia), (Site 15) | Picea abies - Southern Karelian spruce | DeAngelis DL, RH Gardner, HH Shugart. 1981. Productivity of forest ecosystems studies during the IBP: the woodlands data set. In: Dynamic Properties of Forest Ecosystems (DE Reichle, ed) pp. 567-672. Cambridge University Press, Cambridge, UK; Kazimirov NI, RM Morozova. 1973. Biological cycling of matter in spruce forests of Karelia. NAUKA, Leningrad Branch, Academy of Sciences. 168 pp. |
| 54 | Russia (Southern Karelia), (Site 16) | *Picea abies* - Southern Karelian spruce | DeAngelis DL, RH Gardner, HH Shugart. 1981. Productivity of forest ecosystems studies during the IBP: the woodlands data set. In: Dynamic Properties of Forest Ecosystems (DE Reichle, ed) pp. 567-672. Cambridge University Press, Cambridge, UK; Kazimirov NI, RM Morozova. 1973. Biological cycling of matter in spruce forests of Karelia. NAUKA, Leningrad Branch, Academy of Sciences. 168 pp. |
| 55 | Russia (Southern Karelia), (Site 17) | *Picea abies* - Southern Karelian spruce | DeAngelis DL, RH Gardner, HH Shugart. 1981. Productivity of forest ecosystems studies during the IBP: the woodlands data set. In: Dynamic Properties of Forest Ecosystems (DE Reichle, ed) pp. 567-672. Cambridge University Press, Cambridge, UK; Kazimirov NI, RM Morozova. 1973. Biological cycling of matter in spruce forests of Karelia. NAUKA, Leningrad Branch, Academy of Sciences. 168 pp. |
| 56 | Russia (Southern Karelia), (Site 2) | *Picea abies* - Southern Karelian spruce | DeAngelis DL, RH Gardner, HH Shugart. 1981. Productivity of forest ecosystems studies during the IBP: the woodlands data set. In: Dynamic Properties of Forest Ecosystems (DE Reichle, ed) pp. 567-672. Cambridge University Press, Cambridge, UK; Kazimirov NI, RM Morozova. 1973. Biological cycling of matter in spruce forests of Karelia. NAUKA, Leningrad Branch, Academy of Sciences. 168 pp. |
| 57 | Russia (Southern Karelia), (Site 3) | *Picea abies* - Southern Karelian spruce | DeAngelis DL, RH Gardner, HH Shugart. 1981. Productivity of forest ecosystems studies during the IBP: the woodlands data set. In: Dynamic Properties of Forest Ecosystems (DE Reichle, ed) pp. 567-672. Cambridge University Press, Cambridge, UK; Kazimirov NI, RM Morozova. 1973. Biological cycling of matter in spruce forests of Karelia. NAUKA, Leningrad Branch, Academy of Sciences. 168 pp. |
| 58 | Russia (Southern Karelia), (Site 4) | *Picea abies* - Southern Karelian spruce | DeAngelis DL, RH Gardner, HH Shugart. 1981. Productivity of forest ecosystems studies during the IBP: the woodlands data set. In: Dynamic Properties of Forest Ecosystems (DE Reichle, ed) pp. 567-672. Cambridge University Press, Cambridge, UK; Kazimirov NI, RM Morozova. 1973. Biological cycling of matter in spruce forests of Karelia. NAUKA, Leningrad Branch, Academy of Sciences. 168 pp. |
| 59 | Russia (Southern Karelia), (Site 5) | *Picea abies* - Southern Karelian spruce | DeAngelis DL, RH Gardner, HH Shugart. 1981. Productivity of forest ecosystems studies during the IBP: the woodlands data set. In: Dynamic Properties of Forest Ecosystems (DE Reichle, ed) pp. 567-672. Cambridge University Press, Cambridge, UK; Kazimirov NI, RM Morozova. 1973. Biological cycling of matter in spruce forests of Karelia. NAUKA, Leningrad Branch, Academy of Sciences. 168 pp. |
| 60 | Russia (Southern Karelia), (Site 6) | *Picea abies* - Southern Karelian spruce | DeAngelis DL, RH Gardner, HH Shugart. 1981. Productivity of forest ecosystems studies during the IBP: the woodlands data set. In: Dynamic Properties of Forest Ecosystems (DE Reichle, ed) pp. 567-672. Cambridge University Press, Cambridge, UK; Kazimirov NI, RM Morozova. 1973. Biological cycling of matter in spruce forests of Karelia. NAUKA, Leningrad Branch, Academy of Sciences. 168 pp. |
| 61 | Russia (Southern Karelia), (Site 7) | *Picea abies* - Southern Karelian spruce | DeAngelis DL, RH Gardner, HH Shugart. 1981. Productivity of forest ecosystems studies during the IBP: the woodlands data set. In: Dynamic Properties of Forest Ecosystems (DE Reichle, ed) pp. 567-672. Cambridge University Press, Cambridge, UK; Kazimirov NI, RM Morozova. 1973. Biological cycling of matter in spruce forests of Karelia. NAUKA, Leningrad Branch, Academy of Sciences. 168 pp. |
| 62 | Russia (Southern Karelia), (Site 8 | *Picea abies* - Southern Karelian spruce | DeAngelis DL, RH Gardner, HH Shugart. 1981. Productivity of forest ecosystems studies during the IBP: the woodlands data set. In: Dynamic Properties of Forest Ecosystems (DE Reichle, ed) pp. 567-672. Cambridge University Press, Cambridge, UK; Kazimirov NI, RM Morozova. 1973. Biological cycling of matter in spruce forests of Karelia. NAUKA, Leningrad Branch, Academy of Sciences. 168 pp. |
| 63 | Russia (Southern Karelia), (Site 8 | *Picea abies* - Southern Karelian spruce | DeAngelis DL, RH Gardner, HH Shugart. 1981. Productivity of forest ecosystems studies during the IBP: the woodlands data set. In: Dynamic Properties of Forest Ecosystems (DE Reichle, ed) pp. 567-672. Cambridge University Press, Cambridge, UK; Kazimirov NI, RM Morozova. 1973. Biological cycling of matter in spruce forests of Karelia. NAUKA, Leningrad Branch, Academy of Sciences. 168 pp. |
| 64 | Russia (Southern Karelia), (Site 9) | *Picea abies* - Southern Karelian spruce | Gower ST, O Krankina, RJ Olson, M Apps, S Linder, C Wang. 2001. Net primary production and carbon allocation patterns of boreal forest ecosystems. Ecol. Appl 11(5), 1395-1411 |
| 65 | Russia, Karelia | *Picea abies* | Gower ST, O Krankina, RJ Olson, M Apps, S Linder, C Wang. 2001. Net primary production and carbon allocation patterns of boreal forest ecosystems. Ecol. Appl 11(5), 1395-1411 |
| 66 | Russia, Tomsk | *Pinus sylvestris* | Kajimoto T, Y Matsuura, MA Sofronov, AV Volokitina, S Mori, A Osawa, AP Abaimov. 1999. Above- and belowground biomass and net primary productivity of a *Larix gmelinii* stand near Tura, central Siberia. Tree Physiology 19, 815-822; PokrovskyO.S, B Dupré, J Schott. 2005. Fe–Al–organic Colloids Control of Trace Elements in Peat Soil Solutions: Results of Ultrafiltration and Dialysis. Aquat Geochem 11, 241–278; Pokrovsky OS, J Schott, DI Kudryavtzev, B Dupré. 2005. Basalt weathering in Central Siberia under permafrost conditions. Geochim Cosmochim Acta 69:5659–5680; Wang S, L Zhou, J Chen, W Ju, X Feng, W Wu. 2011. Relationships between net primary productivity and stand age for several forest types and their influence on China's carbon balance. Journal of Environmental Management 92: 1651-1662. |
| 67 | Sweden | *Picea abies* | DeAngelis DL, RH Gardner, HH Shugart. 1981. Productivity of forest ecosystems studies during the IBP: the woodlands data set. In: Dynamic Properties of Forest Ecosystems (DE Reichle, ed) pp. 567-672. Cambridge University Press, Cambridge, UK; Gower ST, O Krankina, RJ Olson, M Apps, S Linder, C Wang. 2001. Net primary production and carbon allocation patterns of boreal forest ecosystems. Ecol. Appl 11(5), 1395-1411; Ladanai S. 2008. Nutrient relations in coniferous forests. Doctoral Thesis. Swedish University of Agricultural Sciences. Uppsala 2008. ISBN 978-91-86195-4. <http://pub.epsilon.slu.se/1933/1/kapan.pdf>; Linder S, GI Agren. 2013. NPP Boreal Forest: Jadraas, Sweden, 1973-1990, R1. Data set. Available on-line [http://daac.ornl.gov] from Oak Ridge National Laboratory Distributed Active Archive Center, Oak Ridge, Tennessee, USA. doi:10.3334/ORNLDAAC |
| 68 | Sweden, Jädraås [SWECON] | *Pinus sylvestris* | Gower ST, O Krankina, RJ Olson, M Apps, S Linder, C Wang. 2001. Net primary production and carbon allocation patterns of boreal forest ecosystems. Ecol. Appl 11(5), 1395-1411; Ladanai S. 2008. Nutrient relations in coniferous forests. Doctoral Thesis. Swedish University of Agricultural Sciences. Uppsala 2008. ISBN 978-91-86195-4. <http://pub.epsilon.slu.se/1933/1/kapan.pdf>; Linder S, GI Agren. 2013. NPP Boreal Forest: Jadraas, Sweden, 1973-1990, R1. Data set. Available on-line [http://daac.ornl.gov] from Oak Ridge National Laboratory Distributed Active Archive Center, Oak Ridge, Tennessee, USA. doi:10.3334/ORNLDAAC |
| 69 | Sweden, Jädraås [SWECON] | *Pinus sylvestris* | Bergh J, Freeman M, Sigurdsson B, Kellomäki S, Laitinen K, Niinistö S, Peltola H, Linder S. 2003. Modelling the short-term effects of climate change on the productivity of selected tree species in Nordic countries. Forest Ecol Manage 183, 327-340; Eliasson, P.E. 2007. Impacts of climate change on carbon and nitrogen cycles in boreal forest ecosystems. PhD Dissertation. ISSN: 1652-6880, ISBN: 978-91-576-7388-6; Gower ST, O Krankina, RJ Olson, M Apps, S Linder, C Wang. 2001. Net primary production and carbon allocation patterns of boreal forest ecosystems. Ecol. Appl 11(5), 1395-1411; Linder S. 2013. NPP Boreal Forest: Flakaliden, Sweden, 1986-1996, R1. Data set. Available on-line [http://daac.ornl.gov] from Oak Ridge National Laboratory Distributed Active Archive Center, Oak Ridge, Tennessee, USA. doi:10.3334/ORNLDAAC/201 |
| 70 | Sweden, Northern [Flakaliden] | *Picea abies* | Linder S. 2013. NPP Boreal Forest: Flakaliden, Sweden, 1986-1996, R1. Data set. Available on-line [http://daac.ornl.gov] from Oak Ridge National Laboratory Distributed Active Archive Center, Oak Ridge, Tennessee, USA. doi:10.3334/ORNLDAAC/201 |
| TEMPERATE FORESTS | | | |
| 71 | Slovakia, southern part of the Veporske vrchy massif | *Fagus sylvatica* L. | Konôpka B, J Pajtík, K Noguchi, M Lukac. 2013. Replacing Norway spruce with European beech: A comparison of biomass and net primary production patterns in young stands. Forest Ecology and management 302, 185-192. |
| 72 | Slovakia, southern part of the Veporske vrchy massif | *Picea abies* (L.) H Karst | Konôpka B, J Pajtík, K Noguchi, M Lukac. 2013. Replacing Norway spruce with European beech: A comparison of biomass and net primary production patterns in young stands. Forest Ecology Management 302, 185-192. |
| 73 | Belgium | *Pinus sylvestris* - Belgian Scots pine | Janssens IA, DA Sampson, J Cermák, L Meiresonne, F Riguzzi, S Overloop, R Ceulemans. 1999. Above- and below-ground phytomass and carbon storage in a Belgian Scots pine stand. Ann For Sci 56, 81–90; Yuste Curiel J, B Konôpka, IA Janssens, K Coenen, CW Xiao, R Ceulemans. 2005. Contrasting net primary productivity and carbon distribution between neighboring stands of *Quercus robur* and *Pinus sylvestris*. Tree Physio 25(6), 701-712. |
| 74 | Belgium | *Pinus sylvestris* - Belgian Scots pine | Xiao CW, J Curiel Yuste, IA Janssens, P Roskams, L Nachtergale, A Carrara, BY Sanchez, R Ceulemans. 2003. Above- and below-ground biomass and net primary production in a 73-year-old Scots pine forest. Tree Physiol 23, 505–516 |
| 75 | Belgium | *Quercus robur* | Janssens IA, DA Sampson, J Cermák, L Meiresonne, F Riguzzi,S Overloop, R Ceulemans. 1999. Above- and below-ground phytomass and carbon storage in a Belgian Scots pine stand. Ann For Sci 56, 81–90; Yuste Curiel J, B Konôpka, IA Janssens, K Coenen, CW Xiao, R Ceulemans. 2005. Contrasting net primary productivity and carbon distribution between neighboring stands of *Quercus robur* and *Pinus sylvestris*. Tree Physio 25(6), 701-712. |
| 76 | Belgium | *Quercus* mixed | Cole DW, M Rapp. 1981. Elemental cycling in forests. In: Dynamic Properties of Forest Ecosystems (DE Reichle, ed), pp. 341-409. International Biological Programme 23, Cambridge Univ. Press, London; Vogt, K.A.; Grier, C.C.; Vogt, D.J. 1986. Production, turnover, and nutrient dynamics  of above and belowground detritus of world forests. In: Advances in Ecological Research. Vol. 15. (A MacFadyen, ED Ford, eds), pp. 303-377. London: Academic Press, Harcourt Brace Jovanovich Publishers |
| 77 | Canada, British Columbia | *Pinus contorta*, mesic | Comeau PG, JP Kimmins. 1989. Above- and below-ground biomass and production of Lodgepole pine on sites with differing soil moisture regimes. Can J of Forest Res 19(4), 447-454; Comeau PG, JP Kimmins. 2013. NPP Boreal Forest: Canal Flats, Canada, 1984, R1. Data set. Available on-line [http://daac.ornl.gov] from Oak Ridge National Laboratory Distributed Active Archive Center, Oak Ridge, Tennessee |
| 78 | Canada, British Columbia | *Pinus contorta*, xeric | Comeau PG, JP Kimmins. 1989. Above- and below-ground biomass and production of Lodgepole pine on sites with differing soil moisture regimes. Can J of Forest Res 19(4), 447-454; Comeau PG, JP Kimmins. 2013. NPP Boreal Forest: Canal Flats, Canada, 1984, R1. Data set. Available on-line [http://daac.ornl.gov] from Oak Ridge National Laboratory Distributed Active Archive Center, Oak Ridge, Tennessee |
| 79 | Canada, British Columbia | *Pinus contorta*, xeric | Comeau PG, JP Kimmins. 1989. Above- and below-ground biomass and production of Lodgepole pine on sites with differing soil moisture regimes. Can J of Forest Res 19(4), 447-454; Comeau PG, JP Kimmins. 2013. NPP Boreal Forest: Canal Flats, Canada, 1984, R1. Data set. Available on-line [http://daac.ornl.gov] from Oak Ridge National Laboratory Distributed Active Archive Center, Oak Ridge, Tennessee |
| 80 | Canada, British Columbia [Canal Flats] | *Pinus contorta*, mesic | Comeau PG, JP Kimmins. 1989. Above- and below-ground biomass and production of Lodgepole pine on sites with differing soil moisture regimes. Can J of Forest Res 19(4), 447-454; Comeau PG, JP Kimmins. 2013. NPP Boreal Forest: Canal Flats, Canada, 1984, R1. Data set. Available on-line [http://daac.ornl.gov] from Oak Ridge National Laboratory Distributed Active Archive Center, Oak Ridge, Tennessee |
| 81 | China | *Pinus koraiensis* | Jianping S, T Dali, W Miao, Z Shidong. 1993. Fine roots turnover in a broad-leaved Korean pine forest of Changbai mountain. Chinese J of Applied Ecology 3, 1993. |
| 82 | China - Liangshui National Nature Reserve | *Pinus koraiensis* old-growth | Cai H, X Di, SC Chang, C Wang, B Shi, P Geng, G Jin. 2016. Carbon storage, net primary production, and net ecosystem production in four major temperate forest types in northeastern China. Can J Forest Res 46(2), 143-151; Wang H, X Dong. 2017. Seed plant diversity of different forest types in Liangshui National Natural Reserve. Biodiversity Data Journal 5(10):e22167. DOI: [10.3897/BDJ.5.e22167](https://www.researchgate.net/deref/http%3A%2F%2Fdx.doi.org%2F10.3897%2FBDJ.5.e22167?_sg%5B0%5D=m3k67j_s8cC-N8yu3fWEoZOEki4nE1veQ2VxrUljiEbG8W8k9x5mOjbzWtVR-QzSnN66q00AUqnpZcyYlHBr3f5FOg.a42ML6Tnd-aiD61hvR_A7JTTuBI6So6YcMwraU5gWEi86BUn2BZ3as2i6HBLA8MoEnkappsZdWMC6fDxh4qu6A) |
| 83 | China - Liangshui National Nature Reserve | *Betula platyphylla* - secondary | Cai H, X Di, SC Chang, C Wang, B Shi, P Geng, G Jin. 2016. Carbon storage, net primary production, and net ecosystem production in four major temperate forest types in northeastern China. Can J Forest Res 46(2), 143-151; Wang H, X Dong. 2017. Seed plant diversity of different forest types in Liangshui National Natural Reserve. Biodiversity Data Journal 5(10):e22167. DOI: [10.3897/BDJ.5.e22167](https://www.researchgate.net/deref/http%3A%2F%2Fdx.doi.org%2F10.3897%2FBDJ.5.e22167?_sg%5B0%5D=m3k67j_s8cC-N8yu3fWEoZOEki4nE1veQ2VxrUljiEbG8W8k9x5mOjbzWtVR-QzSnN66q00AUqnpZcyYlHBr3f5FOg.a42ML6Tnd-aiD61hvR_A7JTTuBI6So6YcMwraU5gWEi86BUn2BZ3as2i6HBLA8MoEnkappsZdWMC6fDxh4qu6A) |
| 84 | Colorado, USA | *Picea engelmannii/Abies lasiocarpa* | Arthur MA, TJ Fahey. 1992. Biomass and nutrients in an Engelmann spruce subalpine fir forest in north central Colorado: pools, annual production, and internal cycling. Can J Forest Res. 22(3), 315-325; Arthur MA, TJ Fahey. 1993. Controls on Soil Solution Chemistry in a Subalpine Forest in North‐Central Colorado. Soil Sci Soc Am J 57(4), 1122-1130; Walthall PM. 1985. Acidic deposition and the soil environment of Loch Vale Watershed in Rocky Mountain National Park. Ph.D. Dissertation, Colorado State University. 148 pp. |
| 85 | Denmark | *Fagus sylvatica* | DeAngelis DL, RH Gardner, HH Shugart. 1981. Productivity of forest ecosystems studies during the IBP: the woodlands data set. In: Dynamic Properties of Forest Ecosystems (DE Reichle, ed) pp. 567-672. Cambridge University Press, Cambridge, UK. |
| 86 | England | *Quercus-Betula-Fraxinus* | Cole DW, M Rapp. 1981. Elemental cycling in forests. In: Dynamic Properties of Forest Ecosystems (DE Reichle, ed) pp. 341-409. Cambridge University Press, Cambridge, UK. International Biological Programme 23, Cambridge Univ. Press, London; DeAngelis DL, RH Gardner, HH Shugart. 1981. Productivity of forest ecosystems studies during the IBP: the woodlands data set. In: Dynamic Properties of Forest Ecosystems (DE Reichle, ed) pp. 567-672. Cambridge University Press, Cambridge, UK. |
| 87 | Georgia, USA | *Cornus florida* L*/Acer rubrum* L*/Quercus prinus* L *– mature* mixed hardwood | Cromack K. 1973. Litter production and decomposition in a mixed hardwood watershed and a white pine watershed at Coweeta Hydrologic Station, North Carolina. PhD Dissertation, University of Georgia, Athens; McGinty DT. 1976. Comparative root and soil dynamics on a white pine watershed in the hardwood forest in the Coweeta Basin. PhD Dissertation. University of Georgia, Athen, Georgia; Swank WT, DA Crossley jr. 1988. Forest hydrology and ecology at Coweeta. Ecological Studies 66. Springer. 486 pp. |
| 88 | Germany | *Fagus sylvatica* – European beech | DeAngelis DL, RH Gardner, HH Shugart. 1981. Productivity of forest ecosystems studies during the IBP: the woodlands data set. In: Dynamic Properties of Forest Ecosystems (DE Reichle, ed) pp. 567-672. Cambridge University Press, Cambridge, UK; Vogt, K.A.; Grier, C.C.; Vogt, D.J. 1986. Production, turnover, and nutrient dynamicsof above and belowground detritus of world forests. In: Advances in Ecological Research. Vol. 15. (A MacFadyen, ED Ford, eds), pp. 303-377. London: Academic Press, Harcourt Brace Jovanovich Publishers |
| 89 | Germany, Solling Project [SITE B 3] | *Fagus sylvatica* – European beech | Cole DW, M Rapp. 1981. Elemental cycling in forests. In: Dynamic Properties of Forest Ecosystems (DE Reichle, ed) pp. 341-409. Cambridge University Press, Cambridge, UK. International Biological Programme 23, Cambridge Univ. Press, London. |
| 90 | Germany, Solling Project [Site F 2] | Quercus prinus | Cole DW, M Rapp. 1981. Elemental cycling in forests. In: Dynamic Properties of Forest Ecosystems (DE Reichle, ed) pp. 341-409. Cambridge University Press, Cambridge, UK. International Biological Programme 23, Cambridge Univ. Press, London. |
| 91 | Germany, Solling Project [Site F 3] | Pinus strobus | Cole DW, M Rapp. 1981. Elemental cycling in forests. In: Dynamic Properties of Forest Ecosystems (DE Reichle, ed) pp. 341-409. Cambridge University Press, Cambridge, UK. International Biological Programme 23, Cambridge Univ. Press, London. |
| 92 | Japan, Shigayama | Fagus sylvatica | Kimura M. 1963. [Dynamics of vegetation in relation to soil development in northern Yatsugatake Mountains.](https://www.cabdirect.org/cabdirect/abstract/19630603303) Japanese J. Bot. 18, 255-287. |
| 93 | Japan | *Fagus sylvatica* | DeAngelis DL, RH Gardner, HH Shugart. 1981. Productivity of forest ecosystems studies during the IBP: the woodlands data set. In: Dynamic Properties of Forest Ecosystems (DE Reichle, ed) pp. 567-672. Cambridge University Press, Cambridge, UK. |
| 94 | Japan, Kubotaniyama [JPTF-70 Yusuhara] | *Picea abies* | ORNL IBP-JIBP. ORNL Progress Report 1966-71 OF IBP-JIBP Special Committee in Japanese Science Congress 1966. In: Dynamic Properties of Forest Ecosystems (DE Reichle, ed), 605 p. Cambridge University Press, Cambridge, UK. |
| 95 | Japan, Kyoto [School forest, Ashu] | Evergreen broadleaf forest | ORNL IBP-JIBP. ORNL Progress Report 1966-71 OF IBP-JIBP Special Committee in Japanese Science Congress 1966. In: Dynamic Properties of Forest Ecosystems (DE Reichle, ed), 605 p. Cambridge University Press, Cambridge, UK. |
| 96 | Japan, Aya Research Site LTER | Evergreen broadleaf forest – old-growth | Do TV, T Sato, S Saito, O Kozan, H Yamagawa, D Nagamatsu, N Nishimura, T Manabe. 2015. Effects of micro-topograhies on stand structure and tree species diversity in an old-growth evergreen broad-leaved forest, southwestern Japan. Global ecology and Conservation 4, 185-196; Sato T, Y Kominami, S Saito, K Niiyama, T Manabe, H Tanouchi, N Noma, S Yamamoto. 1999. An introduction to the Aya Research Site, a longterm ecological research site, in a warm temperate evergreen broad-leaved forest ecosystems in southwestern Japan: research topics and design. Bull. Kitakyushu Mus. Nat. His. 18, 57–180; Sato T, Y Kominami, S Saito, K Niiyama, H Tanouchi, D Nagamatsu, H Nomiya. 2010. Temporal dynamics and resilience of fine litterfall in relation to typhoon disturbances over 14 years in an old-growth lucidophyllous forest in southwestern Japan. Plant Ecology 208, 187-198. |
| 97 | Japan, Kyoto prefecture, central Japan | Cool temperate deciduous forest | Tateno R, T Hishi, H Takeda. 2004. Above- and belowground biomass and net primary production in a cool-temperate deciduous forest in relation to topographical changes in soil nitrogen. Forest Ecol Management 193(3), 297-306. |
| 98 | Japan, Takatori-yama [JPTF-71 Yusuhara] | *Abies firma* - True fir | ORNL IBP-JIBP. ORNL Progress Report 1966-71 OF IBP-JIBP Special Committee in Japanese Science Congress 1966. In: Dynamic Properties of Forest Ecosystems (DE Reichle, ed), 605 p. Cambridge University Press, Cambridge, UK. |
| 99 | Massachusetts, USA | *Quercus-Acer* - red oak [*Q rubra*], sugar maple [*A saccharum*] | Aber JD, JM Melillo, KJ Nadelhoffer, CA McClaugherty, J. Pastor. 1985. Fine root turnover in forest ecosystems in relation to quantity and form of nitrogen availability: a comparison of two methods. Oecologica 66(3), 317-321; McClaugherty CA, JD Aber, JM Melillo. 1982. The role of fine roots in the organic matter and nitrogen budgets of two forested ecosystems. Ecology 63(5), 1481-1490; McClaugherty CA, JD Aber, JM Melillo. 1984. Decomposition dynamics of fine roots in forested ecosystems. Oikos 42, 378-386; Nadelhoffer KJ, JD Aber, JM Melillo. 1983. Leaf-litter production and soil organic matter dynamics along a nitrogen-availability gradient in Southern Wisconsin (U.S.A.). Can J Forest Res 13(1), 12-21; Pastor J, JD Aber, CA McClaugherty. 1984. [Aboveground production and N and P cycling along a nitrogen mineralization gradient on Blackhawk Island, Wisconsin](javascript:void(0)). Ecology 65(1), 256-268; Vitousek PM, JR Gosz, CC Grier, JM Melillo, WA Reiners. 1982. A comparative analysis of potential nitrification and nitrate mobility in forest ecosystems. Ecol. Monographs 52(2), 155-177. |
| 100 | Massachusetts, USA | *Pinus resinosa* | Aber JD, JM Melillo, KJ Nadelhoffer, CA McClaugherty, J. Pastor. 1985. Fine root turnover in forest ecosystems in relation to quantity and form of nitrogen availability: a comparison of two methods. Oecologica 66(3), 317-321; McClaugherty CA, JD Aber, JM Melillo. 1982. The role of fine roots in the organic matter and nitrogen budgets of two forested ecosystems. Ecology 63(5), 1481-1490; McClaugherty CA, JD Aber, JM Melillo. 1984. Decomposition dynamics of fine roots in forested ecosystems. Oikos 42, 378-386; Vogt KA, CC Grier, DJ Vogt. 1986. Production, turnover, and nutrient dynamics of above and belowground detritus of world forests. In: MacFadyen, A.; Ford, E.D., eds. Advances in ecological research. Vol. 15. London: Academic Press, Harcourt Brace Jovanovich Publishers: 303-377. |
| 101 | Michigan, USA | *Acer saccharum* -hardwood mix, southern | Hendrick KS, RL Hendrick, R Fogel. 1993. The demography of fine roots in response to patches of water and nitrogen. New Phytologist 125(3), 575-580. |
| 102 | Michigan, USA | *Quercus alba* | Hendrick KS, RL Hendrick, R Fogel. 1993. The demography of fine roots in response to patches of water and nitrogen. New Phytologist 125(3), 575-580. |
| 103 | New Hampshire, USA | *Acer spitcatum*, *Betula lutea*, *Fagus grandifolia* | Bormann FH, GE Likens, TG Siccama, RS Pierce, JS Eaton. 1974. The export of nutrients and recovery of stable conditions following deforestation at Hubbard Brook. Ecol Monographs 44(3), 255-277; Gosz JR, GE Likens, FH Bormann. Nutrient content of litter fall on the Hubbard Brook Experimental Forest, New Hampshire. 1972. Ecology 53(5), 769-784; Safford LO. 1974. Effect of fertilization on biomass and nutrient content of fine roots in a beech-birch-maple stand. Plant Soil 40, 349-363; Siccama TG, FH Bormann, GE Likens. 1970. The Hubbard Brook Ecosystem Study: Productivity, nutrients, and phytosociology of the herbaceous layer. Ecological Monograph 40(4), 389-402. |
| 104 | New Hampshire, USA | Northern hardwoods - mix sugar maple, beech, yellow birch, red maple and white ash | Bormann FH, G Likens. 1979. Pattern and Process in a Forested Ecosystems. Springer- Verlag, New York Inc., 253 pp.; Cole DW, M Rapp. 1981. Elemental cycling in forests. In: Dynamic Properties of Forest Ecosystems (DE Reichle, ed) pp. 341-409. Cambridge University Press, Cambridge, UK. International Biological Programme 23, Cambridge Univ. Press, London; Covington WW. 1976. Forest floor organic matter and nutrient content and leaf fall during secondary succession in Northern Hardwoods. Dissertation. Yale University; Covington WW. 1981. Changes in forest floor organic matter and nutrient content following clearcutting in Northern Hardwoods. Ecology 62(1), 41-48; Fahey TJ, JW Hughes. 1994. Fine root dynamics in a northern hardwood forest ecosystem, Hubbard Brook Ex0perimental Forest, NH. J of Ecology 82(3), 533-548; Harmon ME, WL Silver, B Fasth, H Chen, IC Burke, WJ Parton, SC Hart, WS Currie and Lidet. 2009. Long-term patterns of mass loss during the decomposition of leaf and fine root litter: an intersite comparison. Global Change Biol 15(5): 1320-1338. |
| 105 | New Mexico, Mt Taylor [Cibola National Forest] | *Pseudotsuga menziesii var glauca* -Rocky Mt. mixed conifer, control | Horner JT. 1987. The effects of manipulation of nitrogen and water availability on the polyphenol content of Douglas-fir foliage: Implications for ecosystem theory. Dissertation University of New Mexico, Albuquerque, New Mexico; HornerJD, RG Cates, JR Gosz. 1987. Tannin, nitrogen, and cell wall composition of green vs. senescent Douglas-fir foliage. Oecologia 72, 515-519; Gower ST, KA Vogt, CC Grier. 1992. Carbon dynamics of Rocky Mountain Douglas-fir: Influence of water and nutrient availability. Ecological Monographs 62(1), 43-65; White CS, JR Gosz, JD Horner, DI Moore. 1988. Seasonal, annual, and treatment-induced variation in available nitrogen pools and nitrogen-cycling processes in soils of two Douglas-fir stands. Biology Fertility of Soils 6, 93-99.et al 1988. |
| 106 | New Mexico, Mt Taylor [Cibola National Forest] | *Pseudotsuga menziesii var glauca* -Rocky Mt. mixed conifer, fertilized | Gower ST, KA Vogt, CC Grier. 1992. Carbon dynamics of Rocky Mountain Douglas-fir: Influence of water and nutrient availability. Ecological Monographs 62(1), 43-65. |
| 107 | New Mexico, Mt Taylor [Cibola National Forest] | *Pseudotsuga menziesii var glauca* -Rocky Mt. mixed conifer, irrigation | Gower ST, KA Vogt, CC Grier. 1992. Carbon dynamics of Rocky Mountain Douglas-fir: Influence of water and nutrient availability. Ecological Monographs 62(1), 43-65. |
| 108 | New Mexico, Mt Taylor [Cibola National Forest] | *Pseudotsuga menziesii var glauca* -Rocky Mt. mixed conifer, wood chips added | Gower ST, KA Vogt, CC Grier. 1992. Carbon dynamics of Rocky Mountain Douglas-fir: Influence of water and nutrient availability. Ecological Monographs 62(1), 43-65. |
| 109 | New Mexico, Mt Taylor [Cibola National Forest] | *Pseudotsuga menziesii var glauca* -Rocky Mt. mixed conifer, wood chip/irrigation | Gower ST, KA Vogt, CC Grier. 1992. Carbon dynamics of Rocky Mountain Douglas-fir: Influence of water and nutrient availability. Ecological Monographs 62(1), 43-65. |
| 110 | New York, Brookhaven, USA | *Quercus alba, Q coccinea, Pinus rigida* | Whittaker RH, GM Woodwell. 1968. Dimension and production relations of trees and shrubs in the Brookhaven Forest, New York. J of Ecology 56(1), 1-25 |
| 111 | North Carolina | *Acer rubrum/ Quercus prinus* | DeAngelis DL, RH Gardner, HH Shugart. 1981. Productivity of forest ecosystems studies during the IBP: the woodlands data set. In: Dynamic Properties of Forest Ecosystems (DE Reichle, ed) pp. 567-672. Cambridge University Press, Cambridge, UK. |
| 112 | Oregon | *Tsuga heterophylla/Picea sitchensis* | Grier CC. 1976. Biomass, productivity and nitrogen-phosphorus cycles in hemlock-spruce stands of the central Oregon coast. IN: *Western hemlock management* (WA Atkinson, RJ Zasoski, eds) pp. 71-81. College of Forest Resources, University of Washington, Seattle, WA Contribution No. 34; Cole DW, M Rapp. 1981. Elemental cycling in forests. In: Dynamic Properties of Forest Ecosystems (DE Reichle, ed) pp. 341-409. Cambridge University Press, Cambridge, UK. International Biological Programme 23, Cambridge Univ. Press, London. |
| 113 | Oregon | *Tsuga heterophylla/Picea sitchensis* | Grier CC. 1976. Biomass, productivity and nitrogen-phosphorus cycles in hemlock-spruce stands of the central Oregon coast. IN: *Western hemlock management* (WA Atkinson, RJ Zasoski, eds) pp. 71-81. College of Forest Resources, University of Washington, Seattle, WA Contribution No. 34. |
| 114 | Oregon | *Pinus ponderosa* | Law BE, PE Thornton, J Irvine, PM Anthoni, S Van Tuyl. 2001. Carbon storage and fluxes in ponderosa pine forests at different developmental stages. Global Change Biol. 7, 755-777; Pierce LL, SW Running, J Walker. 1994. Regional-scale relationships of leaf area index to specific leaf area and leaf nitrogen content. Ecol Appl 4, 313-321. |
| 115 | Oregon | *Pinus ponderosa* | Law BE, PE Thornton, J Irvine, PM Anthoni, S Van Tuyl. 2001. Carbon storage and fluxes in ponderosa pine forests at different developmental stages. Global Change Biol. 7, 755-777; Peterson DL, RH Waring. 1994. Overview of the Oregon Transect Ecosystem Research Project. Ecological Applications 4(2), 211-225. |
| 116 | Oregon | *Pseudotsuga menziesii/ Tsuga heterophylla* | Harmon ME, K Bible, MG Ryan, DC Shaw, H Chen, J Klopatek, X Li. 2004. Production, respiration, and overall carbon balance in an old-growth *Pseudotsuga-Tsuga* forest ecosystem. Ecosystems 7, 498-512; Shaw DC, JF Franklin, K Bible, J Klopatek, E Freeman, S Greene, GG Parker. 2004. Ecological setting of the Wind River old-growth forest. Ecosystems 7, 427-439. |
| 117 | Oregon, Cascade Head, coast range [OTTER sites, Site 1] | *Picea sitchensis-Tsuga heterophylla* | Runyon J, RH Waring, SN Goward, JM Welles. 1994. Environmental limits on net primary production and light-use efficiency across the Oregon Transect. Ecol. Applications 4(2), 226-237; Waring RH, JJ Landsberg, M Williams. 1998. Net primary production of forests: a constant fraction of gross primary production? Tree Physiology 18(2), 129-134; Waring RH, B Law, B Bond. 2013. NPP Temperate Forest: OTTER Project Sites, Oregon, USA, 1989-1991, R1. Data set available on-line: https://doi.org/10.3334/ORNLDAAC/472 |
| 118 | Oregon, Eastern High Cascades (Control) [OTTER site, Site 5] | *Ponderosa pine* | Runyon J, RH Waring, SN Goward, JM Welles. 1994. Environmental limits on net primary production and light-use efficiency across the Oregon Transect. Ecol. Applications 4(2), 226-237; Waring RH, JJ Landsberg, M Williams. 1998. Net primary production of forests: a constant fraction of gross primary production? Tree Physiology 18(2), 129-134; Waring RH, B Law, B Bond. 2013. NPP Temperate Forest: OTTER Project Sites, Oregon, USA, 1989-1991, R1. Data set available on-line: https://doi.org/10.3334/ORNLDAAC/472 |
| 119 | Oregon, HJ Andrews Experiimental Forest [Watershed 10] | *Pseudotsuga-Acer-Olystichum* (Cool-moist) | Cole DW, M Rapp. 1981. Elemental cycling in forests. In: Dynamic Properties of Forest Ecosystems (DE Reichle, ed) pp. 341-409. Cambridge University Press, Cambridge, UK. International Biological Programme 23, Cambridge Univ. Press, London; Cromack K Jr, P Sollins, WC Graustein, K Speidel, AW Todd, G Spycher, CY Li, RL Todd. 1979. Calcium oxalate accumulation and soil weathering in mats of the hypogeous fungus *Hysterangium crassum*. Soil Bio Biochemistry 11(5), 463-468; Grier CC, RS Logan. 1977. Old-growth *Pseudotsuga menziesii* communities of a western Oregon watershed: Biomass distribution and production budgets. Ecol Monogr 47(4), 373-400; Sollins P, CC Grier, FM McCorison, K Cromack Jr, R Fogel. 1980. The internal element cycles of an old-growth Douglas-fir ecosystem in western Oregon. Ecol Monogr 50 (3): 261-285. |
| 120 | Oregon, HJ Andrews Experimental Forest [Watershed 10] | *Pseudotsuga-Castganopsis* (Xeric), north slope | Cole DW, M Rapp. 1981. Elemental cycling in forests. In: Dynamic Properties of Forest Ecosystems (DE Reichle, ed) pp. 341-409. Cambridge University Press, Cambridge, UK. International Biological Programme 23, Cambridge Univ. Press, London; Cromack K Jr, P Sollins, WC Graustein, K Speidel, AW Todd, G Spycher, CY Li, RL Todd. 1979. Calcium oxalate accumulation and soil weathering in mats of the hypogeous fungus *Hysterangium crassum*. Soil Bio Biochemistry 11(5), 463-468; Grier CC, RS Logan. 1977. Old-growth *Pseudotsuga menziesii* communities of a western Oregon watershed: Biomass distribution and production budgets. Ecol Monogr 47(4), 373-400; Sollins P, CC Grier, FM McCorison, K Cromack Jr, R Fogel. 1980. The internal element cycles of an old-growth Douglas-fir ecosystem in western Oregon. Ecol Monogr 50 (3): 261-285. |
| 121 | Oregon, HJ Andrews Experimental Forest [Watershed 10] | *Pseudotsuga-Castganopsis* (Xeric), south slope | Cole DW, M Rapp. 1981. Elemental cycling in forests. In: Dynamic Properties of Forest Ecosystems (DE Reichle, ed) pp. 341-409. Cambridge University Press, Cambridge, UK. International Biological Programme 23, Cambridge Univ. Press, London; Cromack K Jr, P Sollins, WC Graustein, K Speidel, AW Todd, G Spycher, CY Li, RL Todd. 1979. Calcium oxalate accumulation and soil weathering in mats of the hypogeous fungus *Hysterangium crassum*. Soil Bio Biochemistry 11(5), 463-468; Grier CC, RS Logan. 1977. Old-growth *Pseudotsuga menziesii* communities of a western Oregon watershed: Biomass distribution and production budgets. Ecol Monogr 47(4), 373-400; Sollins P, CC Grier, FM McCorison, K Cromack Jr, R Fogel. 1980. The internal element cycles of an old-growth Douglas-fir ecosystem in western Oregon. Ecol Monogr 50 (3): 261-285. |
| 122 | Oregon, HJ Andrews Experimental Forest [Watershed 10] | *Pseudotsuga-Rhododenron-Berberis* (Mesic) | Cole DW, M Rapp. 1981. Elemental cycling in forests. In: Dynamic Properties of Forest Ecosystems (DE Reichle, ed) pp. 341-409. Cambridge University Press, Cambridge, UK. International Biological Programme 23, Cambridge Univ. Press, London; Cromack K Jr, P Sollins, WC Graustein, K Speidel, AW Todd, G Spycher, CY Li, RL Todd. 1979. Calcium oxalate accumulation and soil weathering in mats of the hypogeous fungus *Hysterangium crassum*. Soil Bio Biochemistry 11(5), 463-468; Grier CC, RS Logan. 1977. Old-growth *Pseudotsuga menziesii* communities of a western Oregon watershed: Biomass distribution and production budgets. Ecol Monogr 47(4), 373-400; Sollins P, CC Grier, FM McCorison, K Cromack Jr, R Fogel. 1980. The internal element cycles of an old-growth Douglas-fir ecosystem in western Oregon. Ecol Monogr 50 (3): 261-285. |
| 123 | Oregon, Santiam Pass, High Cascades summit [OTTER sites, Site 4] | *Tsuga mertensiana* | Pierce LL, SW Running, J Walker. 1994. Regional-scale relationships of leaf area index to specific leaf area and leaf nitrogen content. Ecol Appl 4(2), 313-321; Runyon J, RH Waring, SN Goward, JM Welles. 1994. Environmental limits on net primary production and light-use efficiency across the Oregon transect. Ecological Applications 4(2), 226-237; Waring RH, B Law, B Bond. 2013. NPP Temperate Forest: OTTER Project Sites, Oregon, USA, 1989-1991, R1. Data set available on-line: https://doi.org/10.3334/ORNLDAAC/472. |
| 124 | Oregon, Scio (control, west Cascades) [OTTER sites, Site 3] | *Tsuga heterophylla-Pseudotsuga menziesii* | Pierce LL, SW Running, J Walker. 1994. Regional-scale relationships of leaf area index to specific leaf area and leaf nitrogen content. Ecol Appl 4(2), 313-321; Runyon J, RH Waring, SN Goward, JM Welles. 1994. Environmental limits on net primary production and light-use efficiency across the Oregon transect. Ecological Applications 4(2), 226-237; Waring RH, B Law, B Bond. 2013. NPP Temperate Forest: OTTER Project Sites, Oregon, USA, 1989-1991, R1. Data set available on-line: https://doi.org/10.3334/ORNLDAAC/472. |
| 125 | Oregon, Waring's Woods, interior valley [OTTER sites, Site 2] | *Pseudotsuga menziesii* | Pierce LL, SW Running, J Walker. 1994. Regional-scale relationships of leaf area index to specific leaf area and leaf nitrogen content. Ecol Appl 4(2), 313-321; Runyon J, RH Waring, SN Goward, JM Welles. 1994. Environmental limits on net primary production and light-use efficiency across the Oregon transect. Ecological Applications 4(2), 226-237; Waring RH, B Law, B Bond. 2013. NPP Temperate Forest: OTTER Project Sites, Oregon, USA, 1989-1991, R1. Data set available on-line: https://doi.org/10.3334/ORNLDAAC/472. |
| 126 | Poland, Ispina, Niepolomice near Krakow | Tilio-Carpinetum, oak-hornbeam forest [*Quercus patraea, Carpinus betulus*] | Bandolo-Ciolczyk E. 1974. Production of tree leaves and energy flow through the litter in Tilio-Carpinetum association (International Biological Programme area). Stud Nat Sci A 9, 29-91. |
| 127 | Russia, Koinas [Arkangelsk Region] | *Picea abies, Juniperus communis, Vaccinium myrtillus* | DeAngelis DL, RH Gardner, HH Shugart. 1981. Productivity of forest ecosystems studies during the IBP: the woodlands data set. In: Dynamic Properties of Forest Ecosystems (DE Reichle, ed) pp. 567-672. Cambridge University Press, Cambridge, UK. |
| 128 | Sweden | *Fagus sylvatica* | DeAngelis DL, RH Gardner, HH Shugart. 1981. Productivity of forest ecosystems studies during the IBP: the woodlands data set. In: Dynamic Properties of Forest Ecosystems (DE Reichle, ed) pp. 567-672. Cambridge University Press, Cambridge, UK. |
| 129 | Sweden | *Fagus sylvatica* | DeAngelis DL, RH Gardner, HH Shugart. 1981. Productivity of forest ecosystems studies during the IBP: the woodlands data set. In: Dynamic Properties of Forest Ecosystems (DE Reichle, ed) pp. 567-672. Cambridge University Press, Cambridge, UK. |
| 130 | Sweden | *Picea abies* | DeAngelis DL, RH Gardner, HH Shugart. 1981. Productivity of forest ecosystems studies during the IBP: the woodlands data set. In: Dynamic Properties of Forest Ecosystems (DE Reichle, ed) pp. 567-672. Cambridge University Press, Cambridge, UK. |
| 131 | Sweden | *Quercus robus-Tilia cordata* | Andersson F. 1970. Ecological studies in a Scanian woodland and meadow area, Southern Sweden. I. Vegetational and environmental structure. Opera Botanica 27, 190 pp.; Andersson F. 1970. Ecological studies in a Scanian woodland and meadow area, Southern Sweden. II. Plant biomass, primary production and turnover of organic matter. Bot Notiser 123, 8-51; Andersson F. 1973. In Modeling Forest Ecosystems. EDFB-IBP 73-7, OAK RIDGE, USA; Nihlgård B. 1969. Microclimate in a beech and a spruce forest; a comparative study from Kongalund, Scania, Sweden. Bot Notiser 122, 333-352; Nihlgård B. 1970. Precipitation, its chemical composition and effect on soil water in a beech and a spruce forest in South Sweden. Oikos 21, 208-17; Nihlgård B. 1971. Pedological influence of spruce planted on former beech forest soils in Scania, South Sweden. Oikos 22, 301-13; Nihlgård B. 1972. Plant biomass, primary production and distribution of chemical elements in a beech and a planted spruce forest in South Sweden. OIKOS 23(1), 69-81. |
| 132 | Sweden [Kongalund Beech Site] | *Fagus sylvatica, Stellaria nemorum* - Beech forest | Andersson F. 1970. Ecological studies in a Scanian woodland and meadow area, Southern Sweden. I. Vegetational and environmental structure. Opera Botanica 27, 190 pp.; Andersson F. 1970. Ecological studies in a Scanian woodland and meadow area, Southern Sweden. II. Plant biomass, primary production and turnover of organic matter. Bot Notiser 123, 8-51; Andersson F. 1973. In Modeling Forest Ecosystems. EDFB-IBP 73-7, OAK RIDGE, USA; Nihlgård B. 1969. Microclimate in a beech and a spruce forest; a comparative study from Kongalund, Scania, Sweden. Bot Notiser 122, 333-352; Nihlgård B. 1970. Precipitation, its chemical composition and effect on soil water in a beech and a spruce forest in South Sweden. Oikos 21, 208-17; Nihlgård B. 1971. Pedological influence of spruce planted on former beech forest soils in Scania, South Sweden. Oikos 22, 301-13; Nihlgård B. 1972. Plant biomass, primary production and distribution of chemical elements in a beech and a planted spruce forest in South Sweden. OIKOS 23(1), 69-81. |
| 133 | Sweden [Kongalund Spruce Site] | *Picea abies* | Nihlgård B, L Lindregn. 1977. Plant biomass, primary production and bioelements of three mature beech forests in South Sweden. Oikos 28, 95-104; Lindgren L. 1970. Beech forest vegetation in Sweden - a survey. Botaniska notiser 123: 401–24. |
| 134 | Sweden, Oved | *Fagus sylvatica* | DeAngelis DL, RH Gardner, HH Shugart. 1981. Productivity of forest ecosystems studies during the IBP: the woodlands data set. In: Dynamic Properties of Forest Ecosystems (DE Reichle, ed) pp. 567-672. Cambridge University Press, Cambridge, UK. |
| 135 | Tennessee, Walker Branch, USA | *Liriodendron tulipifera* | Cole DW, M Rapp. 1981. Elemental cycling in forests. In: Dynamic Properties of Forest Ecosystems (DE Reichle, ed) pp. 341-409. Cambridge University Press, Cambridge, UK. International Biological Programme 23, Cambridge Univ. Press, London; Cox TL, TL Harris, BS Ausmus, NT Edwards. 1978. The role of roots in biogeochemical cycles in an eastern deciduous forest. Pedobiologica 18, 264-271; DeAngelis DL, RH Gardner, HH Shugart. 1981. Productivity of forest ecosystems studies during the IBP: the woodlands data set. In: Dynamic Properties of Forest Ecosystems (DE Reichle, ed) pp. 567-672. Cambridge University Press, Cambridge, UK; Harris WF, RS Kinerson Jr, NT Edwards. 1977. Comparison of below-ground biomass of natural deciduous forest and loblolly pine plantations. Pedobiologia 17, 369-381. |
| 136 | Tennessee, Walker Branch, USA | *Liriodendron tulipifera* | DeAngelis DL, RH Gardner, HH Shugart. 1981. Productivity of forest ecosystems studies during the IBP: the woodlands data set. In: Dynamic Properties of Forest Ecosystems (DE Reichle, ed) pp. 567-672. Cambridge University Press, Cambridge, UK. |
| 137 | Tennessee, Walker Branch, USA | *Pinus echinata* | Climate of the States. 1980. NOAA National Centers for Environmental Information. <https://www.ncdc.noaa.gov/climateatlas/>; Cole DW, M Rapp. 1981. Elemental cycling in forests. In: Dynamic Properties of Forest Ecosystems (DE Reichle, ed), pp. 341-409. International Biological Programme 23, Cambridge Univ. Press, London; DeAngelis DL, RH Gardner, HH Shugart. 1981. Productivity of forest ecosystems studies during the IBP: the woodlands data set. In: Dynamic Properties of Forest Ecosystems (DE Reichle, ed) pp. 567-672. Cambridge University Press, Cambridge, UK. |
| 138 | Tennessee, Walker Branch, USA | *Quercus alba/Carya* spp. | Cole DW, M Rapp. 1981. Elemental cycling in forests. In: Dynamic Properties of Forest Ecosystems (DE Reichle, ed), pp. 341-409. International Biological Programme 23, Cambridge Univ. Press, London; DeAngelis DL, RH Gardner, HH Shugart. 1981. Productivity of forest ecosystems studies during the IBP: the woodlands data set. In: Dynamic Properties of Forest Ecosystems (DE Reichle, ed) pp. 567-672. Cambridge University Press, Cambridge, UK. |
| 139 | Tennessee, Walker Branch, USA | *Quercus prinus* | Cole DW, M Rapp. 1981. Elemental cycling in forests. In: Dynamic Properties of Forest Ecosystems (DE Reichle, ed), pp. 341-409. International Biological Programme 23, Cambridge Univ. Press, London; DeAngelis DL, RH Gardner, HH Shugart. 1981. Productivity of forest ecosystems studies during the IBP: the woodlands data set. In: Dynamic Properties of Forest Ecosystems (DE Reichle, ed) pp. 567-672. Cambridge University Press, Cambridge, UK. |
| 140 | Virginia, USA | *Chamae-cyparis thyoides,* wetland site [Atlantic White Cedar] | Day FP Jr. 1982. Litter decomposition rates in the seasonally flooded Great Dismal Swamp. Ecology 63, 670-678; Day FP. 1984. Biomass and litter accumulation in the Great Dismal Swamp. In, Cypress swamps (KC Ewel, HT Odum, eds.) pp. 386-392. University Presses of Florida, Gainesville; Gomez MM, FP Day Jr. 1982. Litter nutrient content and production in the Great Dismal Swamp. Am. J. Bot. 69, 1314-1321; Megonigal JP, FP Day Jr. 1988. Organic matter dynamics in four seasonally flooded forest communities of the Dismal Swamp. Am. J. Bot. 75, 1334-1343; Powell SW, FP Day Jr. 1991. Root production in four communities in the Great Dismal Swamp. American J of Botany 78(2), 288-297. |
| 141 | Virginia, USA | *Acer rubrum, Nyssa Aquatica* [maple-gum] and oak (*Quercus* spp.) Hardwood mixture Dismal Swamp Upland sites | Day FP Jr. 1982. Litter decomposition rates in the seasonally flooded Great Dismal Swamp. Ecology 63, 670-678; Day FP. 1984. Biomass and litter accumulation in the Great Dismal Swamp. In, Cypress swamps (KC Ewel, HT Odum, eds.) pp. 386-392. University Presses of Florida, Gainesville; Gomez MM, FP Day Jr. 1982. Litter nutrient content and production in the Great Dismal Swamp. Am. J. Bot. 69, 1314-1321; Megonigal JP, FP Day Jr. 1988. Organic matter dynamics in four seasonally flooded forest communities of the Dismal Swamp. Am. J. Bot. 75, 1334-1343; Powell SW, FP Day Jr. 1991. Root production in four communities in the Great Dismal Swamp. American J of Botany 78(2), 288-297. |
| 142 | Virginia, USA | *Nyssa aquatica, Acer rubrum, N. sylvatica* [water tupelo, cottongum, wild olive, large tupelo, sourgum] Wetland site | Day FP Jr. 1982. Litter decomposition rates in the seasonally flooded Great Dismal Swamp. Ecology 63, 670-678; Day FP. 1984. Biomass and litter accumulation in the Great Dismal Swamp. In, Cypress swamps (KC Ewel, HT Odum, eds.) pp. 386-392. University Presses of Florida, Gainesville; Gomez MM, FP Day Jr. 1982. Litter nutrient content and production in the Great Dismal Swamp. Am. J. Bot. 69, 1314-1321; Megonigal JP, FP Day Jr. 1988. Organic matter dynamics in four seasonally flooded forest communities of the Dismal Swamp. Am. J. Bot. 75, 1334-1343; Powell SW, FP Day Jr. 1991. Root production in four communities in the Great Dismal Swamp. American J of Botany 78(2), 288-297. |
| 143 | Virginia, USA | *Taxodium distichum* [Bald cypress] | Day FP Jr. 1982. Litter decomposition rates in the seasonally flooded Great Dismal Swamp. Ecology 63, 670-678; Day FP. 1984. Biomass and litter accumulation in the Great Dismal Swamp. In, Cypress swamps (KC Ewel, HT Odum, eds.) pp. 386-392. University Presses of Florida, Gainesville; Gomez MM, FP Day Jr. 1982. Litter nutrient content and production in the Great Dismal Swamp. Am. J. Bot. 69, 1314-1321; Megonigal JP, FP Day Jr. 1988. Organic matter dynamics in four seasonally flooded forest communities of the Dismal Swamp. Am. J. Bot. 75, 1334-1343; Powell SW, FP Day Jr. 1991. Root production in four communities in the Great Dismal Swamp. American J of Botany 78(2), 288-297. |
| 144 | Washington, USA | *Pseudotsuga menziesii* | Vogt DJ. 1987. Douglas-fir ecosystems in western Washington biomass and production as related to site quality and stand age. PhD Dissertation. University of Washington, Seattle, Washington, USA. |
| 145 | Washington | *Pseudotsuga menziesii* | Vogt DJ. 1987. Douglas-fir ecosystems in western Washington biomass and production as related to site quality and stand age. PhD Dissertation. University of Washington, Seattle, Washington, USA. |
| 146 | Washington, USA | *Pseudotsuga menziesii* | Vogt DJ. 1987. Douglas-fir ecosystems in western Washington biomass and production as related to site quality and stand age. PhD Dissertation. University of Washington, Seattle, Washington, USA. |
| 147 | Washington, USA | *Pseudotsuga menziesii* | Vogt DJ. 1987. Douglas-fir ecosystems in western Washington biomass and production as related to site quality and stand age. PhD Dissertation. University of Washington, Seattle, Washington, USA. |
| 148 | Washington, USA | *Pseudotsuga menziesii* | Vogt DJ. 1987. Douglas-fir ecosystems in western Washington biomass and production as related to site quality and stand age. PhD Dissertation. University of Washington, Seattle, Washington, USA. |
| 149 | Washington, USA | *Pseudotsuga menziesii* | Vogt DJ. 1987. Douglas-fir ecosystems in western Washington biomass and production as related to site quality and stand age. PhD Dissertation. University of Washington, Seattle, Washington, USA. |
| 150 | Washington, USA | *Pseudotsuga menziesii* | Vogt DJ. 1987. Douglas-fir ecosystems in western Washington biomass and production as related to site quality and stand age. PhD Dissertation. University of Washington, Seattle, Washington, USA. |
| 151 | Washington, USA | *Pseudotsuga menziesii* | Vogt DJ. 1987. Douglas-fir ecosystems in western Washington biomass and production as related to site quality and stand age. PhD Dissertation. University of Washington, Seattle, Washington, USA. |
| 152 | Washington, USA | *Pseudotsuga menziesii* | Vogt DJ. 1987. Douglas-fir ecosystems in western Washington biomass and production as related to site quality and stand age. PhD Dissertation. University of Washington, Seattle, Washington, USA. |
| 153 | Washington, USA | *Pseudotsuga menziesii* | Vogt DJ. 1987. Douglas-fir ecosystems in western Washington biomass and production as related to site quality and stand age. PhD Dissertation. University of Washington, Seattle, Washington, USA. |
| 154 | Washington, USA | *Pseudotsuga menziesii* | Vogt DJ. 1987. Douglas-fir ecosystems in western Washington biomass and production as related to site quality and stand age. PhD Dissertation. University of Washington, Seattle, Washington, USA. |
| 155 | Washington, USA | *Pseudotsuga menziesii* | Vogt DJ. 1987. Douglas-fir ecosystems in western Washington biomass and production as related to site quality and stand age. PhD Dissertation. University of Washington, Seattle, Washington, USA. |
| 156 | Washington, USA | *Pseudotsuga menziesii* | Vogt DJ. 1987. Douglas-fir ecosystems in western Washington biomass and production as related to site quality and stand age. PhD Dissertation. University of Washington, Seattle, Washington, USA. |
| 157 | Washington, USA | *Pseudotsuga menziesii* | Vogt DJ. 1987. Douglas-fir ecosystems in western Washington biomass and production as related to site quality and stand age. PhD Dissertation. University of Washington, Seattle, Washington, USA. |
| 158 | Washington, USA | *Pseudotsuga menziesii* | Vogt DJ. 1987. Douglas-fir ecosystems in western Washington biomass and production as related to site quality and stand age. PhD Dissertation. University of Washington, Seattle, Washington, USA. |
| 159 | Washington, USA | *Pseudotsuga menziesii* | Vogt DJ. 1987. Douglas-fir ecosystems in western Washington biomass and production as related to site quality and stand age. PhD Dissertation. University of Washington, Seattle, Washington, USA. |
| 160 | Washington, USA | *Pseudotsuga menziesii* | DeAngelis DL, RH Gardner, HH Shugart. 1991. Productivity of forest ecosystems studies during the IBP: the woodlands data set. In: Dynamic Properties of Forest Ecosystems (Reichle DE, ed) pp. 567-672. Cambridge University Press, Cambridge, UK; Johnson DW, DW Cole, CS Bledsoe, K Cromack Jr, RL Edmonds, SP Gessel, CC Grier, BN Richards, KA Vogt. 1982. Nutrient cycling in forests of the Pacific Northwest. In: Analysis of Coniferous Forest Ecosystems in the Western United States. (RL Edmonds, ed), pp. 186-232. US/IBP Synthesis Series 14. Hutchinson Ross Publishing Co., Pennsylvania, USA; Turner J. 1975. Nutrient cycling in a Douglas-fir ecosystem with respect to age and nutrient status. Unpublished PhD Dissertation, University of Washington, Seattle, Washington, USA. |
| 161 | Washington, USA | *Pseudotsuga menziesii* | Cole DW, SP Gessel. 1968. Cedar River Research. A Program for Studying the Pathways, Rates, and Processes of Elemental Cycling in a Forest Ecosystem. Forest Resources Monograph. Institute of Forest Products, University of Washington College of Forest Resources Contribution No. 4, 54 p.; Vogt KA, DJ Vogt, ST Gower, CC Grier. 1990. Carbon and nitrogen interactions for forest ecosystems. In Above- and below-ground interactions in forest trees in acidified soils. Air Pollution Report 32. (H Persson, ed), pp. 203-235. Commission of the European Communities. Directorate-General for Science, Research and Development. Environment Research Programme, Brussels, Belgium |
| 162 | Washington, USA | *Pseudotsuga menziesii* | Grier CC, JG McColl. 1971. Forest floor characteristics within a small plot in Douglas-fir in Western Washington. Soil Sci Soc Am Proc 35, 988-991; Vogt KA, DJ Vogt, ST Gower, CC Grier. 1990. Carbon and nitrogen interactions for forest ecosystems. In Above- and below-ground interactions in forest trees in acidified soils. Air Pollution Report 32. (H Persson, ed), pp. 203-235. Commission of the European Communities. Directorate-General for Science, Research and Development. Environment Research Programme, Brussels, Belgium |
| 163 | Washington, USA | *Pseudotsuga menziesii,* natural low productivity | Keyes MR, CC Grier. 1981. Below- and above-ground biomass and net production in two contrasting Douglas-fir stands. Can J For Res 11, 599-605. |
| 164 | Washington, USA | *Pseudotsuga menziesii,* natural high productivity | Keyes MR, CC Grier. 1981. Below- and above-ground biomass and net production in two contrasting Douglas-fir stands. Can J Forest Res 11, 599-605. |
| 165 | Washington, USA | Abies amabilis | Grier CC, KA Vogt, MR Keyes, RL Edmonds. 1981. Biomass distribution and above- and belowground production in young and mature *Abies amabilis* zone ecosystems of the Washington Cascades. Can J Forest Res 11, 155-167; Meier C. 1981. The role of fine roots in N and P budgets in young and mature *Abies* *amabilis* ecosystems. PhD Dissertation. University of Washington, Seattle, WA, USA; Vogt KA, CC Grier, CE Meier, RL Edmonds. 1982. Mycorrhizal role in net primary production and nutrient cycling in *Abies amabilis* ecosystems in western Washington. Ecology 63, 370-380; Vogt K. 1991. Carbon budgets of temperate forest ecosystems. Tree Physiology 9, 69-86. |
| 166 | Washington, USA | *Abies amabilis* | Grier CC, KA Vogt, MR Keyes, RL Edmonds. 1981. Biomass distribution and above- and belowground production in young and mature *Abies amabilis* zone ecosystems of the Washington Cascades. Can J Forest Res 11, 155-167; Meier C. 1981. The role of fine roots in N and P budgets in young and mature *Abies* *amabilis* ecosystems. PhD Dissertation. University of Washington, Seattle, WA, USA; Vogt KA, CC Grier, CE Meier, RL Edmonds. 1982. Mycorrhizal role in net primary production and nutrient cycling in *Abies amabilis* ecosystems in western Washington. Ecology 63, 370-380; Vogt K. 1991. Carbon budgets of temperate forest ecosystems. Tree Physiology 9, 69-86. |
| 167 | Wisconsin | *Acer saccharum* | Aber JD, JM Melillo, KJ Nadelhoffer, CA McClaugherty, J Pastor. 1985. Fine root turnover in forest ecosystems in relation to quantity and form of nitrogen availability: a comparison of two methods. Oecologia 66, 317-321; McClaugherty CA, JD Aber, JM Melillo. 1984. Decomposition dynamics of fine roots in forested ecosystems. Oikos 42, 378-386; Nadelhoffer KJ, JD Aber, JM Melillo. 1983. Leaf litter production and soil organic matter dynamics along a nitrogen availability gradient in southern Wisconsin (USA). Can J Forest Res 13, 12-21; Nadelhoffer KJ, JD Aber, JM Melillo. 1985. Fine roots, net primary production and nitrogen availability: a new hypothesis. Ecology 66, 1377-1390. |
| 168 | Wisconsin | *Betula* *alleghaniensis* | Nadelhoffer KJ, JD Aber, JM Melillo. 1983. Leaf litter production and soil organic matter dynamics along a nitrogen availability gradient in southern Wisconsin (USA). Can J Forest Res 13, 12-21; Nadelhoffer KJ, JD Aber, JM Melillo. 1985. Fine roots, net primary production and nitrogen availability: a new hypothesis. Ecology 66, 1377-1390. |
| 169 | Wisconsin | *Quercus rubra* | Aber JD, JM Melillo, KJ Nadelhoffer, CA McClaugherty, J Pastor. 1985. Fine root turnover in forest ecosystems in relation to quantity and form of nitrogen availability: a comparison of two methods. Oecologia 66, 317-321; Nadelhoffer KJ, JD Aber, JM Melillo. 1983. Leaf litter production and soil organic matter dynamics along a nitrogen availability gradient in southern Wisconsin (USA). Can J Forest Res 13, 12-21; Nadelhoffer KJ, JD Aber, JM Melillo. 1985. Fine roots, net primary production and nitrogen availability: a new hypothesis. Ecology 66, 1377-1390. |
| 170 | Wisconsin | *Picea abies* | Nadelhoffer KJ, JD Aber, JM Melillo. 1983. Leaf litter production and soil organic matter dynamics along a nitrogen availability gradient in southern Wisconsin (USA). Can J Forest Res 13, 12-21; Nadelhoffer KJ, JD Aber, JM Melillo. 1985. Fine roots, net primary production and nitrogen availability: a new hypothesis. Ecology 66, 1377-1390. |
